# Leading edge maintenance in migrating cells is an emergent property of branched actin network growth

**DOI:** 10.1101/2020.08.22.262907

**Authors:** Rikki M. Garner, Julie A. Theriot

## Abstract

Animal cell migration is predominantly driven by the coordinated, yet stochastic, polymerization of thousands of nanometer-scale actin filaments across micron-scale cell leading edges. It remains unclear how such inherently noisy processes generate robust cellular behavior. We employed high-speed imaging of migrating neutrophil-like HL-60 cells to explore the fine-scale shape fluctuations that emerge and relax throughout the process of leading edge maintenance. We then developed a minimal stochastic model of the leading edge that reproduces this stable relaxation behavior. Remarkably, we find lamellipodial stability *naturally emerges* from the interplay between branched actin network growth and leading edge shape – with no additional feedback required – based on a synergy between membrane-proximal branching and lateral spreading of filaments. These results thus demonstrate a novel biological noise-suppression mechanism based entirely on system geometry. Furthermore, our model suggests that the Arp2/3-mediated ~70-80° branching angle optimally smooths lamellipodial shape, addressing its long-mysterious conservation from protists to mammals.

**Significance Statement:** All cellular functions are driven by the stochastic dynamics of macromolecules, and thus are subject to biological noise. Here, as a model system for noise-suppression in the context of cell migration, we investigate lamellipodial maintenance – where thousands of stochastically polymerizing filaments self-organize into a highly-stable, micron-scale leading edge. Combining experiment and computational modeling, we (1) establish lamellipodial stability is an *emergent property* of dendritically-branched actin network growth, (2) outline a noise-suppression mechanism based on the *geometry* of lamellipodial actin, and (3) determine the evolutionarily-conserved Arp2/3-mediated ~70-80° branching angle optimally suppresses stochastic fluctuations. Our results not only explain the essential role of Arp2/3-mediated branching in lamellipodial formation, but also address the decades-old question of *why* this specific geometry is so well-conserved.

## Introduction

Cell migration driven by actin polymerization plays an essential role in countless organisms spanning the eukaryotic tree of life^1–3^. Across this broad phylogeny, cells have been observed to form a dizzying array of protrusive actin structures, each exhibiting unique physical and biological properties^4^. In all cases, the fundamental molecular unit of these micron-scale structures is the single actin filament, which polymerizes stochastically by addition of single monomers to push the leading edge membrane forward^5-8^. Higher-order actin structures, and the biological functions they robustly enable, are therefore mediated by the collective action of thousands of stochastically growing filaments^9^. It remains an open question how cells control for – or leverage – this inherent stochasticity to maintain stable leading edge protrusions over length and time scales more than three orders of magnitude larger than the scales of actin monomer addition^10,11^.

Perhaps the archetype of dynamically stable actin structures is the lamellipodium, a flat “leaf-like” protrusion that is ~200 nm tall, up to 100 μm wide, and filled with a dense network of dendritically branched actin filaments^12–15^. Cell types that undergo lamellipodial migration (most notably fish epidermal keratocytes and vertebrate neutrophils) can maintain a single, stable lamellipodium for minutes to hours, allowing the cells to carry out their biological functions^16,17^. For example, in their *in vivo* role as first responders of the innate immune system, neutrophils must undergo persistent migration over millimeter-scale distances to reach sites of inflammation and infection^18,19^. Regardless of the cell type, the origins of this striking stability in the face of stochastic actin filament polymerization remain elusive. It has been widely been assumed for decades that some sort of regulatory or mechanical feedback mechanism must be required for lamellipodial shape stability, with extensive experimental efforts identifying membrane tension^20–26^, plasma membrane curvature-sensing proteins^24,27^, a competition for membrane-associated free monomers^28^, and force-feedback via directional filament branching^29^ as potential contributors. The stability of lamellipodia has also been theoretically proposed to depend on the dendritically branched structure of their actin networks, wherein filaments are oriented at an angle relative to the cell’s direction of migration, allowing growing filament tips to spread out laterally along the leading edge as they polymerize^17,30^. (Although any acute angle would permit spreading, we note that filament orientation in cells has been experimentally observed to be highly stereotyped, averaging ±35° relative to the membrane normal^31,32^ – approximately one half of the highly evolutionarily-conserved ~70° branch angle mediated by the Arp2/3 complex^33–35^.) In contrast to the proposed stabilizing role of spreading, several other features of lamellipodial actin are known to impart nonlinearities on network growth, which might amplify stochastic fluctuations. For instance, dendritic branching is an autocatalytic process which can lead to explosive growth^28,36^. In addition, the growth rates of the actin network are dependent on the velocity of the flexible membrane surface it is pushing, in a manner which imparts hysteresis to the system^22,37^. How spreading might interact with these complexities – and the ultimate consequences for maintenance of a stable leading edge – remains unknown.

Seeking to dissect the origins of lamellipodial stability, we pursued complimentary experimental and computational methodologies. First, we performed high-speed, high-resolution microscopy on migrating human neutrophil-like HL-60 cells to monitor their leading edge shape dynamics. In contrast to the remarkable overall lamellipodial stability observed over minutes, high-speed imaging revealed that the leading edge shape is extremely dynamic at shorter time and length scales, constantly undergoing fine-scale fluctuations around the average cell shape. We determined that these shape fluctuations continually dissipate (thereby enabling long time scale lamellipodial maintenance) in a manner quantitatively consistent with viscous relaxation back to the time-averaged leading edge shape. We next developed a minimal stochastic model of branched actin network growth against a flexible membrane, broadly applicable to a wide variety of cell types, that was able to recapitulate the global leading edge stability and fine-scale fluctuation relaxation behavior observed in cells. Our model suggests that the suppression of stochastic fluctuations is an intrinsic, emergent property of collective actin dynamics at the leading edge, as branched network geometry *alone* is necessary and sufficient to generate lamellipodial stability. Moreover, we find that the evolutionarily-conserved geometry, the ~70° branching angle of the Arp2/3 complex, optimally quells shape fluctuations.

## Results

### Fine-scale leading edge shape fluctuations revealed at high spatiotemporal resolution

Neutrophils form lamellipodia that are intrinsically lamellar, maintaining a thin, locally flat sheet of actin even in the absence of support structures like the substrate surface^15^. Here, we study the migration of neutrophil-like HL-60 cells^38^ within quasi-two-dimensional confinement between a glass coverslip and an agarose pad overlay^39^. In addition to serving as an excellent *in vitro* model for neutrophil surveillance of tissues, this assay allows for easy visualization and quantification of lamellipodial dynamics by restraining the lamellipodium to a single imaging plane. Cells in this type of confinement can migrate persistently, maintaining nearly-constant cell shape, for time scales on the order of minutes to hours^16,40^. In order to capture leading edge dynamics on time scales more relevant to the stochastic growth of individual filaments, we performed high-speed (20 Hz) imaging of migrating HL-60 cells. These experiments revealed dynamic, fine-scale fluctuations around the average leading edge shape (Fig. 1a-c, Movie S1, Fig. S1-2, *Methods*), where local instabilities in the leading edge emerge, grow, and then relax. Notably, these previously-unobserved lamellipodial dynamics are phenotypically distinct from – and almost 100-fold faster than – the oscillatory protrusion-retraction cycles seen in other, slower moving cell types (e.g. fibroblasts)^41–43^.

**Fig. 1:**
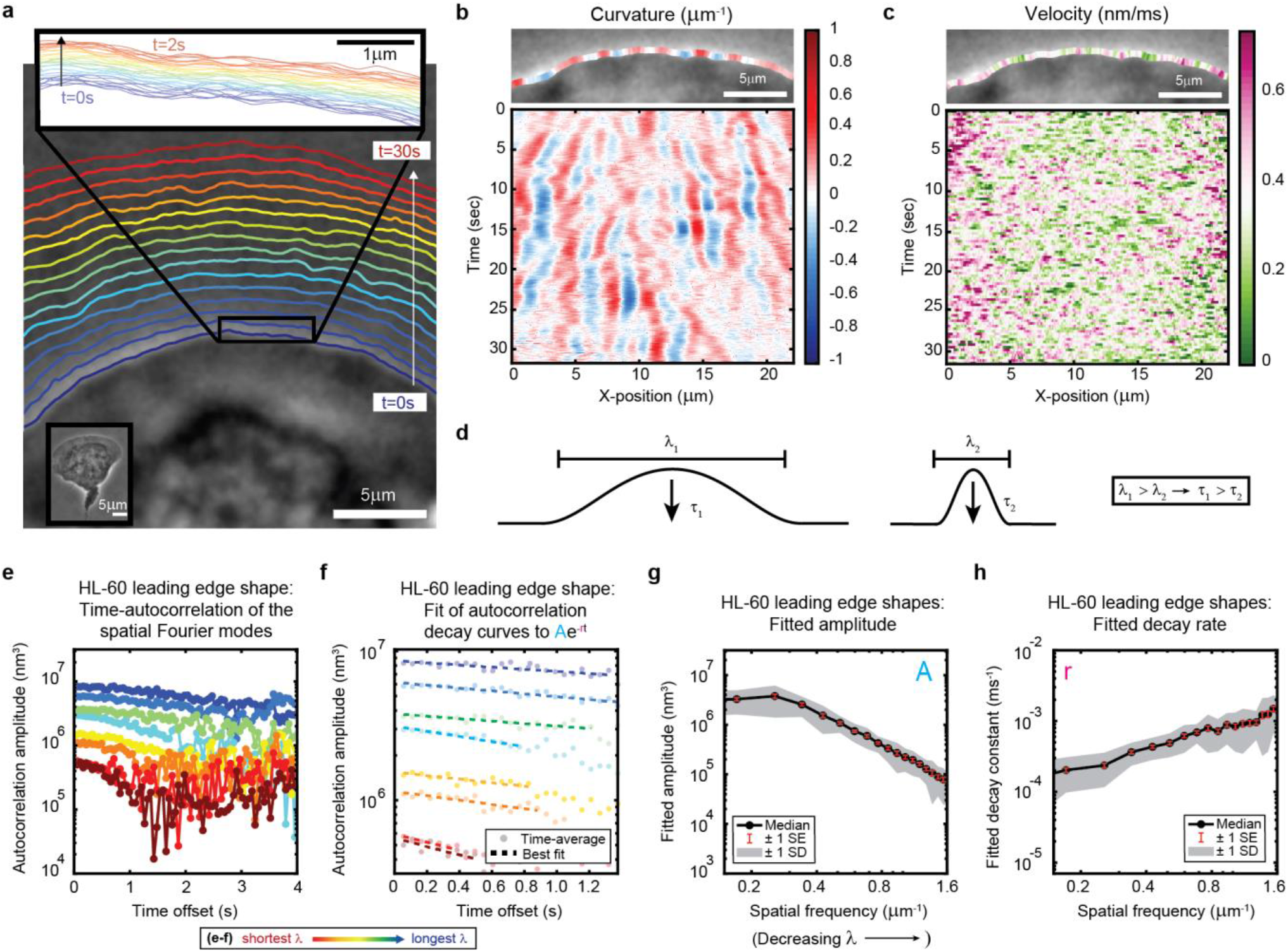
High-speed, high-resolution imaging reveals fine-scale fluctuations in leading edge shape. (**a-c**) Example of leading edge fluctuations extracted from a representative migrating HL-60 cell. (a) Phase contrast microscopy image from the first frame of a movie, overlaid with segmented leading edge shapes from time points increasing from blue to red in 2 sec intervals. Top Inset: Magnification of the segmented leading edge between t = 0-2 sec increasing from blue to red in 50 ms intervals. Bottom Inset: A demagnified image of the whole cell at the last time point. (b-c) Kymographs of curvature (b) and velocity (c). Note the velocity is always positive, so no part of the leading edge undergoes retraction. (**d**) Schematic demonstrating a commonly observed trend between fluctuation wavelength and relaxation time. (**e**) Autocorrelation amplitude (complex magnitude) of the spatial Fourier transform plotted as a function of time offset from a representative cell. Each line corresponds to a different spatial frequency in the range of 0.22-0.62 μm^-1^ (corresponding to a wavelength in the range of 4.5-1.6 μm) in 0.056 μm^-1^ intervals. (**f**) Best fit of the autocorrelation data shown in (e) to an exponential decay, fitted out to a drop in amplitude of 2/*e*. (**g-h**) Fitted parameters of the autocorrelation averaged over 67 cells.

We estimate we were able to reliably measure fluctuations with wavelengths as small as ~650 nm, and amplitudes down to ~65 nm, by fitting the phase contrast halo around the leading edge (Fig. S1-2, *Methods*). These values should approximately correspond to 25 actin filaments at physiological spacing^12^ and 25 actin monomers assembled into a filament lattice along the direction of motion. While our measurements of shape dynamics cannot resolve polymerization events of individual filaments, our results are consistent with the hypothesis that stochasticity in actin growth at the level of monomer addition – occurring throughout the leading edge actin network – ultimately manifests as the observed micron-scale leading edge fluctuations. In particular, kymograph analysis of curvature and velocity (Fig. 1b-c) showed that relatively long-lived shape fluctuations are formed by the continual time-integration of seemingly uncorrelated and very short-lived (sub-second, sub-micron) velocity fluctuations. Because the average cell shape remains constant over time, there must be some form of feedback acting on leading edge curvature to sustain stable lamellipodial growth. These rich, measurable fine-scale dynamics therefore provide a unique opportunity to directly observe the time-evolution of leading edge maintenance. Taking advantage of our high-precision measurements, we aimed to quantitatively investigate the properties of the observed fluctuations, with the goal of determining the mechanisms by which molecular machinery at the leading edge coordinates the stochastic polymerization of individual actin filaments.

### Lamellipodial stability mediated by viscous relaxation of shape fluctuations

The relaxation of fine-scale shape fluctuations back to the steady-state leading edge shape is essential for the long time scale stability of lamellipodia. As for any physical system, the nature of this relaxation reflects the system’s underlying physical properties; in this case, the characteristics of – and interactions between – actin filaments and the membrane. To provide a framework for exploration of the physical mechanisms underlying stable lamellipodial protrusion, we quantified the relaxation dynamics by performing time-autocorrelation analysis on the leading edge shape (*Methods*). Applied in this context, this analytical technique calculates the extent to which the lamellipodium contour loses similarity with the shape at previous time points as fluctuations emerge and relax. As most material systems (actively-driven or otherwise) exhibit relaxation behavior with a characteristic wavelength-dependence (e.g., Fig. 1d), we performed Fourier decomposition on the leading edge shape to separate out fluctuations at different length scales, and then performed autocorrelation analysis separately on each Fourier mode. We validated our analytical methods using simulations of membrane dynamics, for which there exists a well-established analytical theory (*Methods*, Fig. S3), and show that our results are not sensitive to an extension of our analysis to longer length and time scales (*Methods*, Fig. S4-5). Further, the membrane simulation control nicely demonstrates how visual features of curvature kymographs (e.g., Fig. 1b) can be misleading (Methods, Fig. S3), and motivates the necessity of our more comprehensive technique.

Autocorrelation analysis revealed a monotonic relaxation of shape fluctuations at each wavelength (Fig. 1e-f); the decay at every spatial scale is well-fit by an exponential form (Fig. 1f), consistent with overdamped viscous relaxation. Importantly, we do not detect any increase in the autocorrelation over time, which would have appeared if there were any sustained, correlated growth of the fluctuations before they decay. This again suggests that the fluctuations arise from uncorrelated stochastic processes, such as fluctuations in actin density. A clear wavelength-dependence is observed, with shorter wavelengths decaying faster and having smaller amplitudes (Fig. 1g-h). This general trend is shared by many physical systems with linear elastic constraints, such as idealized membranes^44^ and polymers^45^ freely fluctuating under Brownian motion, but can be contrasted with systems that have a dominant wavelength, as in the case of buckling or wrinkling of materials under compression^46^. Importantly, these qualitative and quantitative properties of the leading edge fluctuations are not specific to cell type or experimental conditions (e.g., agarose overlay, ECM), as we also observe this phenomenon in fish epidermal keratocytes (Fig. S6, Movie S2).

### Leading edge stability as an emergent property of branched actin growth

The rich behavior and quantitative nature of our leading edge shape fluctuation data made them ideal for comparison with physical models. In order to understand how molecular-scale actin assembly and biomechanics might give rise to the observed micron-scale shape dynamics, we aimed to reproduce this behavior in a stochastic model of branched actin network growth against a membrane (Fig. 2a-c, Movie S3, *Methods*). Previous stochastic models of protrusive actin-based forces largely focused on actin polymerization against rigid obstacles (e.g., the bacterial cell wall for the *Listeria* comet tail^36,47^ or a single, flat membrane segment in models of lamellipodia^22^). Expanding on this general framework, and in an approach conceptually similar to previous work simulating small (< 2 μm) patches of a lamellipodium^48,49^, we incorporated a two-dimensional leading edge with filaments polymerizing against a flexible membrane, which we modeled as a system of flat membrane segments coupled elastically to each other. The size of the membrane segments was comparable to the spatial resolution of our experimental measurements, allowing us to assay fluctuations over a similar dynamic range of wavelengths. Simulated filaments apply force to the membrane following the classic untethered Brownian ratchet formalism^5^, consistent with recent experiments showing that cellular protrusions are formed by largely untethered actin networks^50^. Designed to be as comparable as possible to our experimental data, the model incorporated experimentally-measured values from the literature for the membrane tension, membrane bending modulus, and biochemical rate constants (Table 1-2)^5,51^. As we were specifically interested in identifying biophysical mechanisms regulating leading edge stability, we minimized the model’s biological complexity by including only the core biochemical elements of actin network growth dynamics: polymerization, depolymerization, branching, and capping. All filament nucleation in the model occurs through dendritic branching (observed in cells to be mediated by the Arp2/3 complex^3,12^, which catalyzes the nucleation of new “daughter” actin filaments as branches from the sides of pre-existing “mother” filaments at a characteristic angle of ~70° ^33–35^). By simulating individual filament kinetics, the model captures the evolutionary dynamics of the filament network, allowing us to directly test hypothesized mechanisms for the interplay between actin network properties (e.g., filament orientation) and protrusion dynamics^5,17,22,30,31,48^ (Fig. 2b).

**Fig. 2:**
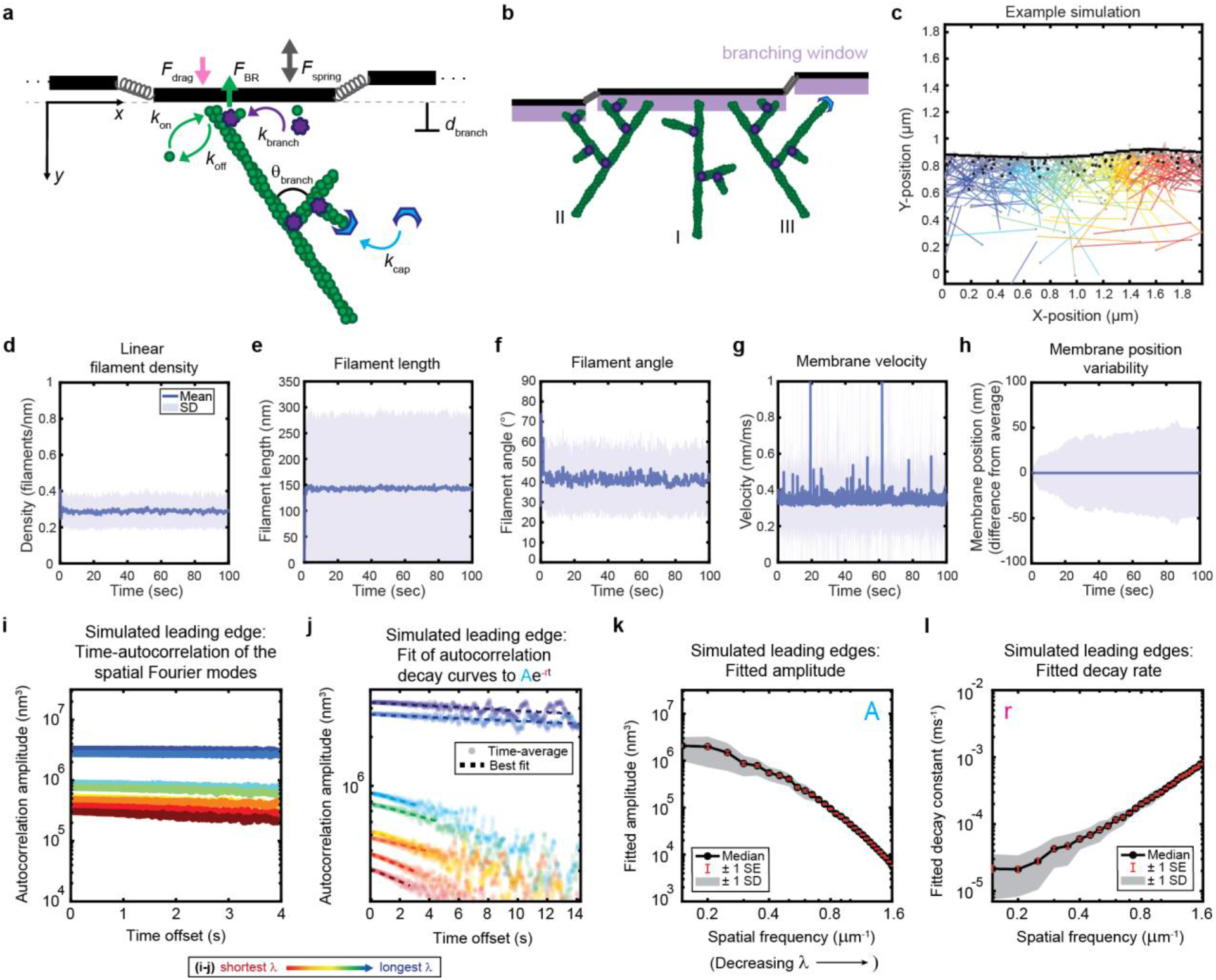
Minimal model of branched actin growth recapitulates leading edge stability and shape fluctuation relaxation. (**a-b**) Model schematic. Black lines, membrane; green circles, actin; purple flowers, Arp2/3 complex; blue crescents, capping protein. Rates: *k*_on_, polymerization; *k*_off_, depolymerization; *k*_branch_, branching; *k*_cap_, capping. *θ*_branch_, branching angle. *d*_branch_, branching window. Physical parameters: *F*_spring_, forces between membrane segments; *F*_BR_, force of filaments on the membrane (Brownian ratchet); *F*_drag_, viscous drag. (b) Schematic demonstrating filament angle evolution. Filaments growing perpendicular to the leading edge (I) outcompete their progeny (branches), leading to a reduction in filament density; filaments growing at an angle (II and III) make successful progeny. Filaments spreading down a membrane positional gradient (II) are more evolutionarily successful than those spreading up (III). (**c**) Simulation snapshot: Black lines, membrane; colored lines, filament equilibrium position and shape; gray dots, barbed ends; black dots, capped ends; filament color, x-position of membrane segment filament is pushing (increasing across the x-axis from blue to red). (**d-h**) For a representative simulation, mean (solid line) and standard deviation (shading) of various membrane and actin filament properties as a function of simulation time. Note for linear filament density (d) lamellipodia are ~10 filament stacks tall along the z-axis, giving mean filament spacing of 10/density ~ 30nm. (**i-l**) Autocorrelation analysis and fitting for a representative simulation (i-j) as well as best fit parameters averaged over 40 simulations (k-l).

**Table 1.** Actin network growth parameters. Parameters listed are the default used for the simulations.

| Notation | Meaning | Value | Source |
| --- | --- | --- | --- |
| $M$ | Free monomer concentration | 15 $\mu\text{M}$ | 76,77 |
| $k_{\text{on}}$ | Polymerization rate | $11 \cdot 10^{-3} \text{ monomers ms}^{-1} \mu\text{M}^{-1}$ | 78 |
| $k_{\text{off}}$ | Depolymerization rate | $10^{-3} \text{ monomers ms}^{-1}$ | 78 |
| $k_{\text{cap}}$ | Capping rate | $3 \cdot 10^{-3} \text{ ms}^{-1}$ | $\sim 3 \cdot 10^{-3} \mu\text{M}^{-1} \text{ ms}^{-1}$ 79<br>at 1 $\mu\text{M}$ capping protein 80 |
| $k_{\text{branch}}$ | Branching rate | $4.5 \cdot 10^{-5} \text{ branches ms}^{-1} \mu\text{M}^{-1} \text{ nm}^{-1}$ | 50nm branch spacing 12,58;<br>Branch rate approximated such that<br>elongation rate / branch rate = 50nm;<br>$k_{\text{branch}} = (k_{\text{on}} \cdot M \cdot l_{\text{m}}) / (50\text{nm} \cdot M \cdot y_{\text{branch}})$ |
| $y_{\text{branch}}$ | Branching window length | 15 nm | $\sim 3\text{-}5$ protein diameters<br>away from the membrane |
| $\theta_{\text{branch}}$ | Branching angle | $70 \pm 10^\circ$ | 33–35,56–58 |
| $l_{\text{p}}$ | Actin filament persistence length | 1 $\mu\text{m}$ | 81 |
| $l_{\text{m}}$ | Actin monomer length | 2.7 nm | 78 |

**Table 2.**
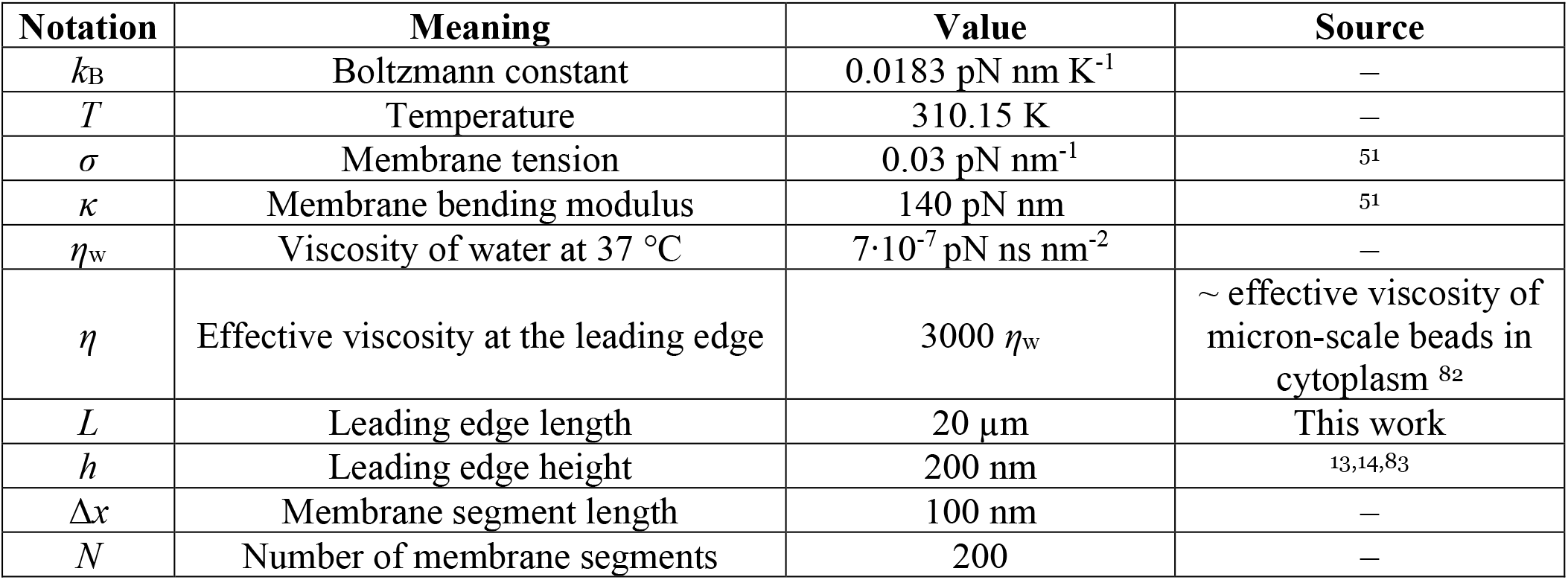
Physical parameters. Parameters listed are the default used for the simulations.

| Notation | Meaning | Value | Source |
| --- | --- | --- | --- |
| $k_{\text{B}}$ | Boltzmann constant | 0.0183 pN nm K<sup>-1</sup> | — |
| $T$ | Temperature | 310.15 K | — |
| $\sigma$ | Membrane tension | 0.03 pN nm<sup>-1</sup> | 51 |
| $\kappa$ | Membrane bending modulus | 140 pN nm | 51 |
| $\eta_{\text{w}}$ | Viscosity of water at 37 °C | $7 \cdot 10^{-7} \text{ pN ns nm}^{-2}$ | — |
| $\eta$ | Effective viscosity at the leading edge | $3000 \eta_{\text{w}}$ | ∼ effective viscosity of<br>micron-scale beads in<br>cytoplasm 82 |
| $L$ | Leading edge length | 20 $\mu\text{m}$ | This work |
| $h$ | Leading edge height | 200 nm | 13,14,83 |
| $\Delta x$ | Membrane segment length | 100 nm | — |
| $N$ | Number of membrane segments | 200 | — |

To our great surprise, this very simple model was able to recapitulate stable leading edge fluctuations. Nascent leading edges reach steady state values for filament density, filament length, filament angle, membrane velocity, and (most importantly) membrane fluctuation amplitude within seconds, a biologically realistic time scale (Fig. 2d-h). Furthermore, the steady state values obtained are in quantitative agreement with both our own experimental data and previously published measurements, with the model yielding mean values of: 0.3 filaments/nm for filament density (~30 nm filament spacing for a lamellipodium that is 10 filaments tall^13^), ~150 nm for filament length, ~40° for filament angle (with respect to the direction of migration), ~0.35 nm/ms for membrane velocity, and ~50 nm for membrane fluctuation amplitude^12,31,32^.

As observed in the experimental measurements, simulated leading edge shape stability is mediated by an exponential decay of shape fluctuations (Fig. 2i-j). Furthermore, the minimal model correctly predicts the monotonic trends of fluctuation amplitude and decay time scale with wavelength (Fig. 2k-l) in a way that was not sensitive to our choices of simulation time step, membrane segment length, and overall length of the leading edge (Fig. S7-9). It should be noted that the generation of the simulated data in Figure 2k-l did not involve any curve-fitting (and therefore *no free parameters* that could be fit) to the experimentally-measured autocorrelation dynamics in order to parameterize the model. Rather, the simulated fluctuation relaxation behavior, qualitatively reproducing our experimental measurements, emerges directly from the molecular-scale actin growth model, in which all biochemical parameters were estimated from measurements in the existing literature (Table 1-2) – leaving no free simulation parameters.

### Predicting effects of drug treatment with Latrunculin B

We were interested in further assaying the predictive power of this minimal stochastic model by determining whether the output of the simulations was congruent with experimental observations under conditions that had not been tested prior to model development. As an example, we elected to test whether the model could correctly predict the response of HL-60 cells to treatment with the drug Latrunculin B, which binds to and sequesters actin monomers. Qualitatively, cells treated with Latrunculin B (Movie S4) present with enhanced bleb formation and more variable leading edge shapes, in comparison with cells treated with a DMSO vehicle control (Movie S5). In our model, addition of this drug can be simulated by reducing the free monomer concentration, which consequently reduces both the polymerization rate and the branching rate. At low effective doses, subtle but measurable changes to leading edge fluctuations were predicted: specifically, an increase in the amplitudes at large wavelengths, and a decrease in the decay rates across all wavelengths (Fig 3a-b). Our experimental results were consistent with these quantiative predictions; Latrunculin B-treated cells exhibited increased fluctuation amplitudes and decreased fluctuation rates over the predicted ranges (Fig. 3c-d).

**Fig. 3:**
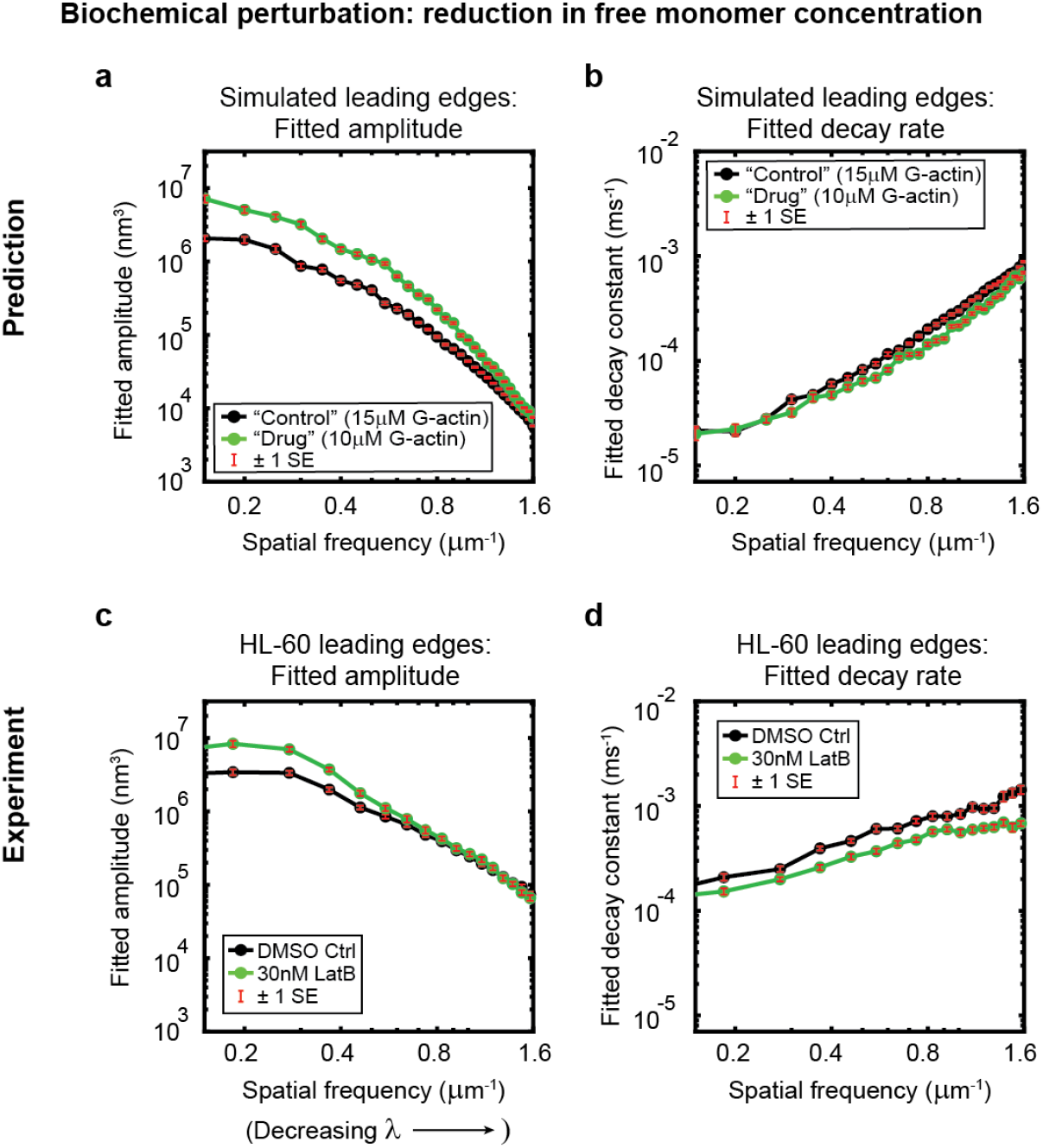
Minimal model correctly predicts response of HL-60 cells to drug treatment. Predicted and experimentally-measured response of the autocorrelation decay fit parameters to drug treatment with Latrunculin B, plotted as in Fig. 1g-h. (**a-b**) Predicted response to a reduction in the free monomer concentration (green, 10 μM G-actin) compared to the standard concentration used in this work (black, 15 μM G-actin) – medians over 40 simulations for each condition. (**c-d**) Experimentally measured behavior: DMSO control – medians over 67 cells (same data as plotted in Fig. 1g-h). 30nM Latrunculin B – medians over 34 cells.

### Geometry as the core determinant of simulated leading edge stability

Given the success of the model in reproducing experimental results, we next wanted to determine which features of the simulation were responsible for leading edge stability and relaxation of fluctuations. The simplicity of the model allowed us to determine the stability mechanism by process of elimination, selectively removing elements of the model (*Methods*) and determining whether stability was retained. To assay the importance of membrane tension and bending rigidity, which has been suggested to be a key factor regulating lamellipodial organization^25,26^, we simply removed the forces between the membrane segments (Fig. 2a, *F_spring_*) from the simulation (*Methods*). Surprisingly, the coupling between the membrane segments (i.e., the effects of tension and bending at length scales larger than the size of an individual membrane segment) was completely dispensable for leading edge stability (Fig. 4a-e).

**Fig. 4:**
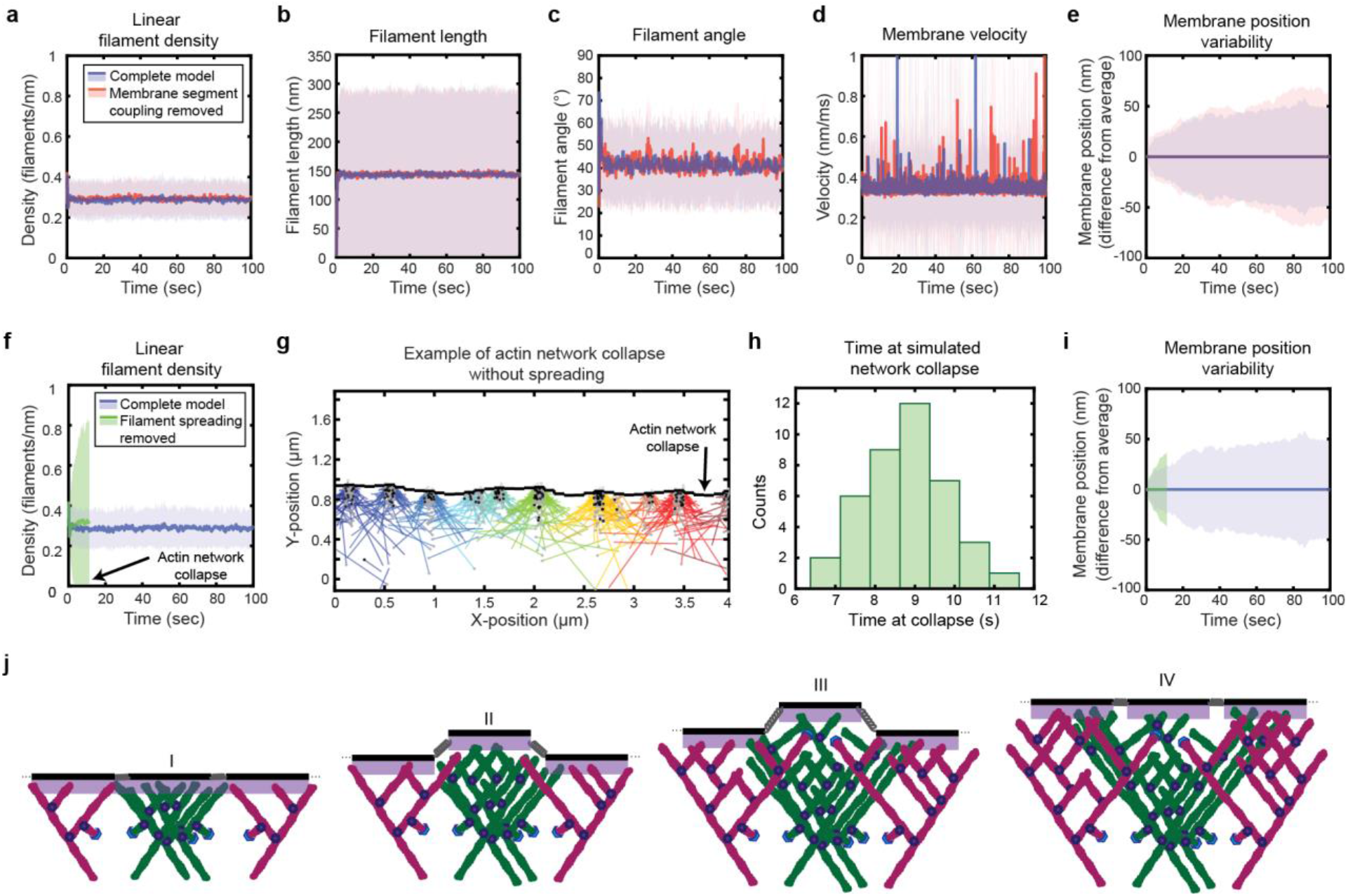
Simulated lamellipodial stability is governed by leading edge geometry. (**a-e**) Comparison of leading edge properties with and without the coupling of the membrane segments by tension and bending rigidity (no coupling: *F_spring_* = 0 in Fig. 2a), plotted as in Fig. 2d-h. (**f-i**) Comparison of leading edge properties with and without the ability of filaments to spread between neighboring membrane segments. (g) A snapshot of the simulation after filament network collapse (defined as a state where at least 25% of the membrane segments have no associated filaments). (f,g,i) Plots made from the same simulation. (h) A histogram of the average time to network collapse over 40 simulations. (**j**) Schematic representing the proposed molecular mechanism underlying the stability of leading edge shape, with time increasing from I-IV.

Following a similar process of elimination, we determined that in fact only two elements were required for stability. First, as reported previously for dendritic actin network polymerization against a single stiff obstacle, it was necessary to constrain branching to occur only within a fixed distance from the leading edge membrane (Fig. 2a-b) in order to maintain a steady state actin density^36^. The molecular motivation for this spatially limited “branching window” is rooted in that fact that activators of the Arp2/3 complex, which render Arp2/3 competent for actin filament nucleation, are typically membrane-associated proteins^52^. Second, we found that stability is inherently tied to the ability of filaments to spread laterally to neighboring membrane segments (Fig. 2b II-III, Fig. 4f-i). (Recall that, because the branched actin network geometry causes filaments to grow, on average, at an angle relative to the membrane normal^31,32^, polymerizing tips spread laterally along the leading edge^17,30^.) Removing filament spreading from the model by fixing filaments to remain associated with their nearest membrane segment at birth (*Methods*) led to actin density divergence: network regions with low filament density eventually underwent complete depolymerization, while high-density regions continued to accumulate actin (Fig. 4f-g).

These findings lead us to a simple molecular feedback mechanism for leading edge stability, based on a synergy between filament spreading and membrane-proximal branching (Fig. 4j): To begin with, regions with initially high filament density come to protrude beyond the average position of the rest of the membrane, representing the emergence of a leading edge shape fluctuation (Fig. 4j, I-II). This induces asymmetric filament spreading, where filaments from high-density regions can spread productively into neighboring regions (Fig. 2b, II), but filaments spreading from adjacent low-density regions cannot keep up with the fast moving membrane segments (Fig. 2b, III), and thus are unproductive (Fig. 4j, II-III). In this way, the branched geometry inherent to dendritic actin polymerization, as well as its interaction with the shape of the membrane, naturally encodes leading edge stability (Fig. 4j, IV). Thus our results directly demonstrate a “stability-through-spreading” mechanism that has previously only been assumed in mean-field analytical theories^17,30^. Remarkably, this means that leading edge maintenance is an intrinsic, emergent property of branched actin network growth against a membrane, without requiring any further regulatory governance. Geometrical constraints imposed simply by the nature of membrane-proximal actin branching ensure that any small variations in *either* local actin filament density *or* growth rate are inherently self-correcting to regress toward the mean.

Of note, it has previously been shown that Arp2/3-mediated branching is required for lamellipodial formation in a wide variety of cell types; cells with inhibited or depleted Arp2/3 complex exhibit complete disruption of the lamellipodium shape and often switch to a different mode of migration altogether, such as filopodial motility^15,53-55^. Indeed, HL-60s treated with the Arp2/3 inhibitor CK-666 have extremely variable leading edge shapes, characterized by long, thin filopodia-like protrusions (Movie S6). Our theoretical results provide a mechanistic interpretation for this striking phenomenon, suggesting that the vital lamellipodial maintenance role of Arp2/3-mediated branching stems from its ability to mediate efficient filament spreading and equilibration of actin density fluctuations, purely because the daughter filament always grows at an angle distinct from its mother.

### Optimal suppression of fluctuations by the highly-conserved ~70° branching angle

Given the essential contribution of branched network geometry to the stability of the simulated leading edges, we reasoned that variations in the branching geometry alone might have a significant effect on leading edge fluctuations. We therefore performed simulations to determine the effects of changing the average branching angle and branching angle variability on filament orientation, filament density, and leading edge fluctuation fit parameters (Fig. 5). In this context, we highlight the distinction between the branching angle, *θ_br_* (i.e., the angle of a daughter filament relative to its mother), and the filament angle or orientation, *θ_f_* (i.e., the angle of a filament relative to the direction of migration) (Fig. 5a, inset). Due to the sterotypical branching angle, *θ_br_*, there is a direct correspondence between the orientation, *θ_f_^mother^*, of a mother filament and the orientation, *θ_f_^daughter^*, she passes on to her daughter branches. Our simulations are thus, in effect, selection assays, as mother filaments compete to stay within the fixed branching window, spawn daughter branches, and thus pass down their angle to their progeny^31,48^. For example, when filaments are initialized with a random orientation, and the branching angle is fixed (i.e., there is no variability in the branching angle), only a handful of the initial filament angles (*θ_f_*) survive until the end of the simulation (Fig. 5a). The surviving, successful filament angles are narrowly and symmetrically distributed around one half of the branching angle (Fig. 5a-c). This optimal filament angle allows mother and daughter filaments to branch back and forth symmetrically about the membrane normal, such that mother filaments do not out-compete their progeny (as has been described previously)^31,48,49^ (Fig. 2b).

**Fig. 5:**
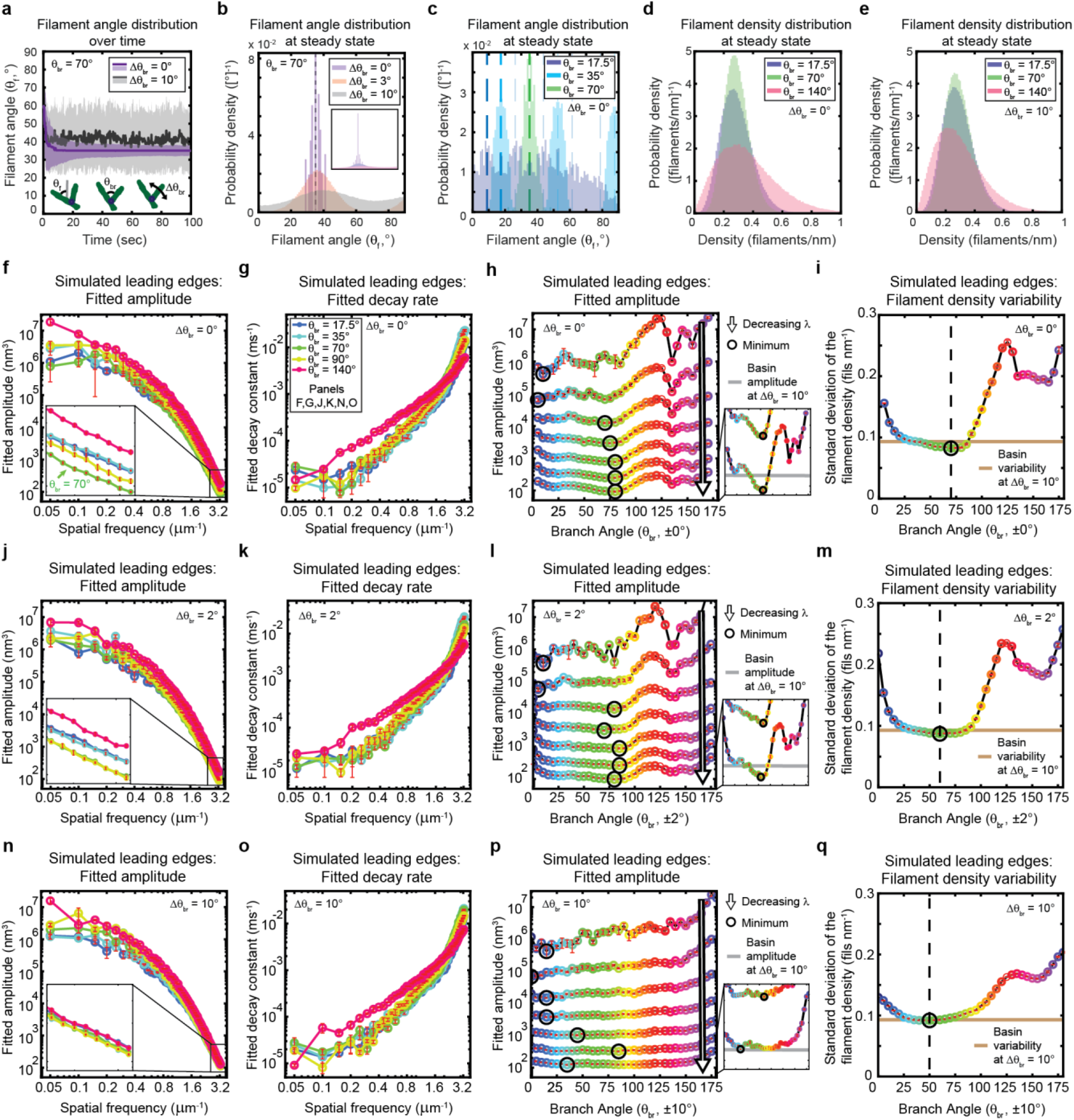
The genetically-encoded Arp2/3-mediated branching angle is optimal for suppressing leading edge fluctuations. (**a-c**) Time course (a) and steady state distribution (b-c) of the filament angle (θ_f_) for simulations with various branching angle standard deviations (Δθ_br_, a-b) and means (θ_br_, c). Dashed lines represent θ_br_/2 plus integer multiples of θ_br_. (**d-e**) Steady state filament density distribution as a function of the mean branching angle in the context of Δθ_br_=0° (d) and Δθ_br_=10° (e). (a-b, d-e) Results from a representative simulation for each condition. (c) Data integrated over 40 simulations. (**f-q**) Predicted response of leading edge fluctuations and filament density variability to changes in the branch angle and branch angle variability, medians over 40 simulations for each condition. Red error bars – standard error. Color map is identical for panels c-q. (h,l,p) Fitted amplitude as a function of branch angle, where each line represents a different spatial frequency, increasing from 0.2-3.2 um^-1^ in intervals of 0.5 um^-1^. Insets have identical x-axes to main panels. (i,m,q) Standard deviation of the filament density at steady state plotted as a function of branch angle. Note the x- and y-axes limits in (f-g, j-k, n-o) are expanded compared to the equivalent panels in Fig. 1-3.

In living cells, branching is mediated by the Arp2/3 complex, which has been experimentally measured to form highly regular and sterotyped branches at ~70° ^33–35^. Intriguingly, this protein complex is highly conserved^3^, with measurements of the branching angle in a wide variety of species (protists^33,35-56^, yeast^35^, mammals^35,56,57^, and amphibians^58^) using various experimental techniques (platinum replica electron microscopy, cryo-electron microscopy, and total internal reflection microscopy) all falling within the range of 67-78° (± 2-13°). The high degree of conservation hints that this specific angle might carry some functional optimality, but the question has not been addressable experimentally; due to the lack of naturally occuring Arp2/3 variants with a substantially different branching angle, an alternative branching structure would hypothetically have to be designed do novo, presumably by altering the protein interaction interface by which the Arp2/3 complex binds to the side of a mother filament^34^. We thus sought to explore the possible functional significance of this conserved angle using our minimal stochastic model. Excitingly, we found that in simulations with no branch angle variability, a 70-80° branching angle was optimal for minimizing both actin density fluctuations (Fig. 5d,i) and leading edge fluctuation amplitudes for wavelengths smaller than ~2 μm (Fig. 5f,h). These smoothing effects are therefore predicted to be relevant within the experimentally-measurable range of wavelengths (between ~0.7 μm and ~2 μm), but are most beneficial for the smallest wavelengths resolved by our simulations (down to ~0.3 μm) – closest to the length scales of individual filament polymerization. Overall, these results provide tantalizing mechanistic insight into the long-standing question of why the characteristic branching angle is so ubiquitous.

Heritability of filament orientation (i.e., the extent to which mother filament orientation determines the orientation of the daughters) is set by the degree of variability in the branching angle (which, in turn, reflects the influence of thermal fluctuations). Perhaps unsurprisingly, decreasing this orientational heritability significantly reduces the dependence of fluctuation amplitude and filament density variability on the branching angle (Fig. 5j-q), and thereby counteracts the beneficial effect of the optimal angle on leading edge fluctuations. Introducing a branch angle variability of ±2° (on the lower end of the experimentally-measured values) broadens the range of near-optimal branch angles but maintains the optimum at ~70-80° (Fig. 5j,l), while introducing a variability of ±10° (on the higher end of the measured range) completely removes the optimum (Fig. 5n,p). In both cases, increasing the branch angle variability increases the minimum possible fluctuation amplitude (Fig. 5h,l,p – insets) and filament density variability (Fig. 5i,m,q), representing a decrease in the noise-suppression capabilities of the system. Overall, these results provide strong support for the idea that actin network geometry is not only essential for leading edge stability, but also plays a major role in determining the fundamental limits of smoothness in lamellipodial shape.

## Discussion

The emergence of robust collective behaviors from stochastic elements is an enduring biological mystery which we are only beginning to unravel^59–65^. The apparent dichotomy in actin-based motility between the random elongation of individual filaments and the stable formation of smooth and persistent higher-order actin structures such as lamellipodia exemplifies this enigma, and provides an avenue towards understanding general strategies for noise suppression in biological systems.

In recent years it has become clear that perturbation-free experiments which examine fluctuations around the mean at steady state (in contrast to probing the change in the mean due to a perturbation) can be a powerful tool for understanding noisy systems^66^. In this work, application of that principle in combination with high-precision measurements, quantitative analytical techniques, and physical modeling led to the surprising revelation that the suppression of stochastic fluctuations *naturally emerges* from the interactions between a growing actin network and the leading edge membrane, with *no additional feedback required*. Our insights into the molecular mechanisms mediating lamellipodial stability were largely enabled by experimentally measuring micron-scale leading edge shape dynamics and comparing them to a molecular-scale actin network growth model that correctly predicts this emergent behavior (as well as many other experimentally-measured features of lamellipodial actin networks). Ultimately, we hope our results inspire future experimental work to directly measure the nanometer-scale interactions predicted by our simulations, which may be accomplished using super-resolution imaging of actin dynamics *in vivo* or in an *in vitro* reconstitution of lamellipodial protrusion (i.e., branched polymerization of actin driving the motion of a flexible barrier).

The model developed in this work provides an understanding of the basic biophysical mechanisms underlying lamellipodial migration. Of course, living cells are home to array of additional complexities which are likely to further modulate leading edge fluctuations and stability, and our model may provide a framework for future exploration of such effects across diverse cell types and experimental conditions. Cells migrating *in vivo* inevitably experience a much more challenging and dynamic environment, in which additional feedback mechanisms will almost certainly be required for the maintenance of polarized migration. The simplicity and biophysical realism of our modeling framework should make it particularly well-suited for future studies focused on predicting and understanding the effects of additional potential feedback mechanisms, including tethering^67–70^, a limiting monomer concentration^28^, force-dependent branching^29,37,71^, and regulation by curvature-sensing proteins^24,72^. Further, this computational model might be useful for exploring the effects of various extracellular forces, such as those produced by obstacles or variations in matrix density, or intracellular forces, such as those produced by hydrostatics. We also note that a certain degree of biochemical signaling is implicit in our model in the form of biochemical rate constants that are invariant in time and space; this property relies on signaling networks to maintain uniform gradients of actin-associated molecules^73^. How local biochemical control (or lack thereof) over these rate constants might affect leading edge fluctuations remains an interesting avenue for future investigation, both theoretically and experimentally.

The defining characteristic and major advance of our model was the explicit inclusion of both the evolutionary dynamics of the actin network and its interaction with the two-dimensional geometry of the leading edge. By selectively removing elements of the model, we determined that lateral filament spreading, combined with a fixed branching window, is indispensable for leading edge stability. This highlights the crucial role in lamellipodial maintenance of the branched structure of actin networks, wherein each daughter filament inherits angular information from its mother. Our further investigations into the evolutionary properties of actin network growth revealed that a ~70-80° branch angle maximally suppresses fine-scale actin density and leading edge shape fluctuations, showing for the first time that Arp2/3-mediated branching imparts optimal functionality, as was long hypothesized based on strong sequence, structural, and functional conservation throughout the eukaryotic tree of life. It is interesting to note that the evolutionarily-conserved branching angle that we find maximally suppresses leading edge fluctuations appears not to be the same angle that optimizes the polymerization velocity of single filaments – predicted to be a broad angle closer to ~90-100° (and quite load-dependent) in the low-load regime^5^. This contrast suggests that evolutionary selection acts at the level of actin network properties, rather than force production by individual filaments.

Returning to the broader question of how noisy biological systems control for stochasticity, we find that stability in the case of lamellipodial dynamics is inherently encoded by the geometry of branched actin network growth. It will be interesting to see whether similar principles hold for other cytoskeletal structures with clear geometric constraints, such as endocytic pits, the cytokinetic ring, and the mitotic spindle.

## Materials and Methods

### HL-60 cell culture and differentiation

HL-60 cells were cultured as described previously^39,40^. In brief, cells were maintained at a density of 0.1-1 x 10^5^ cells/mL by passaging every 2-3 days into fresh RPMI media supplemented with 10% heat-inactivated fetal bovine serum and antibiotics/antimycotics. Supplementation with 1.57% DMSO was used to differentiate the cells into a neutrophil-like state. Cells were subsequently extracted for experiments at 6 days post-differentiation. Our HL-60 cell line was obtained from Orion Weiner’s lab at UCSF, who originally received them from Henry Bourne’s lab at UCSF. The identity of this suspension cell line was confirmed based on the behavior of the cells, including differentiating into a neutrophil-like state upon exposure to DMSO that exhibits characteristic phenotypes for substrate adhesion, rapid migration, and elongated morphology. HL-60s are not listed as a misidentified cell line on the Register of Misidentified Cell Lines. The cell line tested negative to mycoplasma contamination.

### Under-agarose motility assays with HL-60s

Differentiated HL-60 cells were plated on fibronectin-coated coverslips and then overlaid with a 1% agarose pad containing 1nM fMLP (to enhance migratory behavior), as described previously^40^. Microscopy of the migrating cells was performed at 37 °C, using transmitted light to image phase contrast on an epifluorescence microscope at 100X magnification (100x 1.45NA Plan Apo oil objective, Nikon MRD31905). A more detailed description of our microscopy system can be found in previous publications^40^. For treatment with Latrunculin B or CK-666, the drug was embedded into the agarose pad by adding the drug to the unpolymerized agarose pad solution before gelling (at the same time as adding fMLP), such that drug treatment begins when cells are overlaid with the agarose pad and is maintained throughout imaging. Cells were imaged at 45 minutes post-plating. Drugs were first diluted down to 1000X in DMSO, then added to the agarose solution at a dilution of 1:1000 (for a final concentration of 30 nM for Latrunculin B and 100 μM for CK-666), giving a final DMSO concentration in the pad of 0.1%. Controls were performed by adding 0.1% DMSO to the agarose pad alone.

### Keratocyte isolation and motility assays

Keratocytes were cultured from wild-type zebrafish embryos at two days post-fertilization as described previously^74^. Briefly, zebrafish embryos were collected at two days post-fertilization, dechorionated, and anesthetized using tricaine. To dissociate the keratocytes, dechorionated fish were then washed in PBS, incubated in Cell Dissociation Buffer for 30 min at 4°C, incubated in a solution of 0.25% trypsin and 1mM EDTA for ~15 min at 28°C, and then incubated in fetal bovine serum to quench the trypsin. From this point on, cells were maintained in antibiotic and antimycotic to deter microbial growth. The keratocyte-rich supernatant was then concentrated by centrifugation at 500g for 3 min. Keratocytes were then plated on collagen-coated coverslips and incubated at room temperature for ~1 hr to allow cells to adhere. Once adherent, the supernatant was exchanged for imaging media (10% fetal bovine serum in L-15) and allowed to incubate another 15 min at room temperature before imaging. The keratocytes were imaged at 28 °C under similar conditions as those used for HL-60 cells. Data included in Fig. S6 represent cells from a single coverslip. Experiments were approved by University of Washington Institutional Animal Care and Use Committee (protocol 4427-01).

### Image segmentation

Most segmentation algorithms penalize curvature in the contour in order to reduce the noise in the fitting algorithm^75^. However, this runs the risk of introducing artificial correlations and structure into the data. For example, a spring-like curvature penalty would artificially make fluctuations appear to be stretch-dominated. Therefore, we performed the following custom segmentation algorithm to avoid these potential artifacts (Fig. S1). Phase contrast time-lapse videos were manually aligned to the direction of motion of the cell, such that each cell migrates up the y-axis on an x-y coordinate system (Fig. S1a-b). The videos were then cropped to isolate the cell leading edges and exclude the cell body, for easier segmentation. If the cell migrates up the y-axis, this means every image pixel along the x-axis has an associated leading edge position along the y-axis. A leading edge y-position was assigned to each x-axis pixel independently of knowledge about neighboring x-axis pixels, to avoid injecting the artifacts discussed above. A manual segmentation was performed for the first time point in each movie. A custom, automated segmentation algorithm written in MATLAB then performed a line scan of the phase contrast intensity along the y-axis separately for each pixel along the x-axis. For each line scan, the algorithm performs a local search for the leading edge position, constrained to be within a fixed number of pixels from either the manual segmentation (for the first time point) or the previous time point (for subsequent timepoints). The leading edge position is defined as the midpoint between the brightest phase intensity (phase halo) and the point of steepest intensity gradient (transition from phase halo to phase-dense cytoplasm).

### Preparation of the curvature and velocity kymographs

The curvature and velocity kymographs (Fig. 1b-c, Fig. S3) were prepared using custom MATLAB code. Curvatures are calculated as the inverse radius of the best-fit circle corresponding to a 30 pixel-wide (~1.5 μm) region about each position. The most prominent fluctuation events seen in the curvature kymographs (Fig. 1b, Fig. S3) somewhat correspond to (but do not exactly match) the length of the fitting window. For example, the simulated and experimental data shown in Fig. S3 were fit using the same ~1.5 μm fitting window and have similar apparent “dominant wavemodes” of ~3 μm, or twice the fitting window. However, despite being fit with the same fitting window, the simulated data has an observably smaller apparent “dominant wavemode” than the experimental data. Velocities were calculated as the distance traveled over 250 ms (five 50 ms timepoints) non-overlapping windows.

### Processing of segmented cell shapes for autocorrelation analysis

Kymographs of curvature and velocity such as those shown in Fig. 1b-c, while helpful to obtain a qualitative sense of the fluctuation data, are visually dominated by the largest size-scale features of the leading edge. They thus offer an incomplete description of the shape fluctuations^43^ – notably de-emphasizing the fine-scale features that are the subject of this study. Further, curvature kymographs emphasize features that are approximately the same size as the fitting window, and fail to pick up fluctuations at different size scales. To perform a quantitative analysis which faithfully captures fluctuations at all size scales, we choose to perform Fourier decomposition on the leading edge shape, and analyze the dynamics of each wavemode separately. As cells migrate, their global leading edge shape undergoes long timescale changes, such as variations in width, large-scale curvature, or slight turning of the cell, which can dominate the Fourier amplitudes and the subsequent autocorrelation signal. As we are most interested in extracting the fine-scale fluctuations, we performed background subtraction on the segmented leading edge shapes (Fig. S2). To do this, we defined the “background” leading edge shape as the contour after smoothing (by the lowess method, using a span of 7 μm). This rather large smoothing window was chosen specifically to preserve fine-scale features. The background-subtracted y-position is thus defined as the difference between the segmented leading edge and its smoothed counterpart. This process removed the large-scale features of the leading edge. We next wanted to remove the long-timescale features of the leading edge, so we also subtracted the time-averaged background-subtracted y-position for each x-pixel. Altogether, these pre-processing steps maintained the features of interest in the curvature kymograph (Fig. S3). After performing background subtraction, we still needed to control for changes in leading edge width over time. The wavelengths represented in the Fourier transform are defined as *λ*=*L/n*, where *L* is the length of the leading edge, and n is an integer from 0 to one half the number of pixels. If the leading edge length were to vary over time, then so would the wavelengths, making it impossible to track the behavior of a single wavelength fluctuation over time. We thus cropped the dataset along the x-axis to include only pixels which contain the cell for all timepoints in the video, thereby extracting a fixed-length leading edge subset for further analyses.

### Autocorrelation analysis and fitting

Autocorrelation analysis and fitting were performed separately for each cell and simulation. For experimental data, the entire video was analyzed. For simulated data, analysis was only performed on the time points after the simulation had reached steady state, for which we used a conservative cut-off of 10 sec (see Fig. 2d-h). To separate out fluctuations at different length scales, we first performed a spatial Fourier transform on the leading edge shape. Referencing the coordinate system defined in Fig. 2a, the pixels (experiments) or membrane segments (simulations) are equally spaced in the x-direction, allowing us to perform a one-dimensional Fourier transform (MATLAB fft() function, which assumes periodic boundary conditions) on the y-positions of a segmented leading edge for each time point. We then normalize the Fourier transform by a factor of 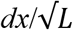, where *dx* is the pixel/membrane segment size and *L* is leading edge length. This normalization preserves the variance and accounts for the pixel size. To measure the fluctuation relaxation, we calculated the time-averaged autocorrelation (*A_n_*(*τ*) = <*Y_n_*(*t*+*τ*)·*Y_n_*\*(*t*)>*_t_*, using non-overlapping windows in *t*) of each Fourier mode amplitude. The autocorrelation function extracted from this analysis contains complex elements of the form A = a + ib. We performed all plotting and fitting on the complex magnitude (sqrt(a^2^ + b^2^)) of the autocorrelation function, which is most representative of the total autocorrelation.

Note that because we are plotting the complex magnitude (which is always positive), the autocorrelation plots shown in Fig. 1e-f and Fig. 2i-j are expected to decay to some non-zero background noise window, rather than to zero. Indeed, the membrane simulation control (see *Validation of autocorrelation analysis implementation*), which is predicted to have a purely exponentially-decaying autocorrelation function, also shows a decay to a noise window at long times (Fig. S4). We fit each Fourier mode time-autocorrelation to the exponential decay function described in the main text (Fig. 1f), fitting *ln*(|*A_n_*(*τ*)|) vs *τ* to a line using MATLAB’s polyfit function, to extract fit parameters for each cell and simulation. Each curve was fit out to a drop in the amplitude by a factor of *e*/2, or at least 10 points. To average fit parameters over many cells, we controlled for cell-to-cell variability in leading edge length by binning the parameters by spatial frequency, and then calculating summary statistics separately for each bin.

The spatial background subtraction performed on the leading edge shape (discussed in Methods: *Processing of segmented cell shapes for autocorrelation analysis*) was necessary to extract the fine-scale shape fluctuations studied in this work. This background subtraction is expected to remove fluctuations with wavelengths larger than ~7 μm (i.e., reduce their amplitude to zero). For this reason, only wavelengths less than 7 μm are plotted in Fig. 1e-h. We note that it is possible that fluctuations with wavelengths less than, but near 7 μm might also have slightly reduced measured amplitudes (i.e., the shape of the curve in Fig. 1g may artificially level off at low spatial frequency). However, any such effect would be performed uniformly in time, and therefore is not expected to affect the measured temporal dynamics (Fig. 1h). Indeed, when we extend the span of our background subtraction by ~50% (up to 10 μm), we find that only the amplitude of the largest mode is altered (slightly increased) and the measured relaxation timescales are not affected (Fig. S5).

### Validation of autocorrelation analysis implementation

To validate our autocorrelation method, we analyzed control simulations of a membrane freely fluctuating under Brownian motion in the absence of actin, and showed it recapitulates predictions from analytical theory for this system (Fig. S3). These simulations were performed exactly as in the leading edge simulations, using the same parameters, but without actin. The equation of motion used for membrane segments in the control simulations, as well as a derivation of the associated autocorrelation function, can be found in Appendix 2.4.4: Choice of timestep. Interestingly, the curvature kymographs of these control simulations exhibit striking visual features reminiscent of instabilities, dominant wavemodes, or oscillations – and yet such effects are absent from this system by definition (which we confirm quantitatively using our autocorrelation method, Fig. S3). This suggests that similar features in the experimentally-measured curvature kymograph are also not indicative of instabilities, dominant wavemodes, or oscillations, which we confirmed by autocorrelation analysis.

### Modeling

Here we briefly describe the geometry and major elements of the model. Please see the Appendix for a detailed description of the model and Tables S1-S2 for a list of the chosen parameters. A simulated patch of leading edge was modeled by a branched network of actin filaments stochastically polymerizing towards a 2D strip of membrane, subject to periodic boundary conditions. The 2D strip was discretized as membrane segments that are fixed in position along one axis and move only along the direction of motion of the simulated cell. Stochastic, fixed time step Brownian dynamics simulations, implemented with custom MATLAB code, were performed to update the membrane position and actin network properties. Actin network growth evolved from constant rate Poisson processes for polymerization, depolymerization, branching, and capping. Once polymerized, filaments were fixed in position at their branch point of origin (in the lab frame of reference), and did not undergo retrograde flow (i.e. translation of the filament position opposite the direction of migration) or translational diffusion. The membrane strip was subject to forces of bending and stretching, drag from fluid viscosity, as well as the force of actin^5,6,36^. Filaments apply force to the membrane segments according to the untethered Brownian ratchet formalism^5^, in which filament pointed end positions are assumed to be rigidly connected to the network (via their branch point of origin) and their barbed end positions are able to freely fluctuate. (As previously, we ignore the possibility of filament buckling due to the fact that lamellipodial filaments exist in a sufficiently low-load, high branch density regime^5^, and experimental evidence shows no indication of buckling^12^.) We expanded this formalism (which previously only considered filament fluctuations perpendicular to the filament’s long axis) to include all fluctuations of the filament along the filament’s short and long axes. Each filament pushes the membrane segment that spans the growing tip’s x-position. We note that the filament angles used to determine the filament forces on the membrane and presented throughout this work are always calculated relative to the global average direction of motion of the leading edge, rather than the local average membrane normal (a simplifying approximation necessitated by the discrete geometry and motivated by the shallow curvatures exhibited by cell leading edges). Control simulations were run to verify that the leading edge fluctuation behavior described in this work was not dependent on the temporal discretization (i.e., simulations were run with sufficiently small timesteps to resolve the fastest dynamics, Fig. S7), spatial discretization (i.e., simulations were run with sufficiently short membrane segments to resolve the smallest length scales at which there is significant bending, Fig. S8), or leading edge length (i.e., the periodic boundary conditions were implemented correctly, such that a simulated small patch of leading edge behaves identically to an equivalently sized portion of a larger simulated patch of leading edge, Fig. S9). In cases where membrane tension and bending rigidity were removed, these forces were simply not calculated in the simulation (Fig. 7a-e). To remove filament spreading, we modified how the filament position was updated upon addition of a monomer in order to maintain the growing filament tip’s x-position. Addition of monomers contributed only to changes in the barbed-end y-position, leaving the x-position intact, while updating the filament length correctly (effectively sliding the pointed end x-position backwards, rather than advancing the barbed end x-position forwards, Fig. 7f-i).

## Acknowledgements

We dedicate this work to A. B. Savinov, who was in preparation alongside this manuscript and was born during revisions. We are exceedingly grateful to E. Labuz for generously isolating and preparing fish epidermal keratocytes used for the experiments presented in Fig. S6. We also kindly thank E. F. Koslover and A. J. Spakowitz for their thoughtful advice on model conceptualization and data analysis, and E. F. Koslover, A. Mogilner, N. M. Belliveau, P. Radhakrishnan, and A. Savinov for helpful comments on the manuscript. Funding: Howard Hughes Medical Institute to J.A.T., Washington Research Foundation to J.A.T., Gerald J. Lieberman Fellowship to R.M.G, NSF Graduate Research Fellowship to R.M.G.

## Data and Materials Availability

Analysis and modeling code for this paper is available on the Theriot lab Gitlab: <https://gitlab.com/theriot_lab/leading-edge-stability-in-motile-cells-is-an-emergent-property-of-branched-actin-network-growth>. All data will be made available upon reasonable request to the corresponding author.

---

**Fig. S1:**
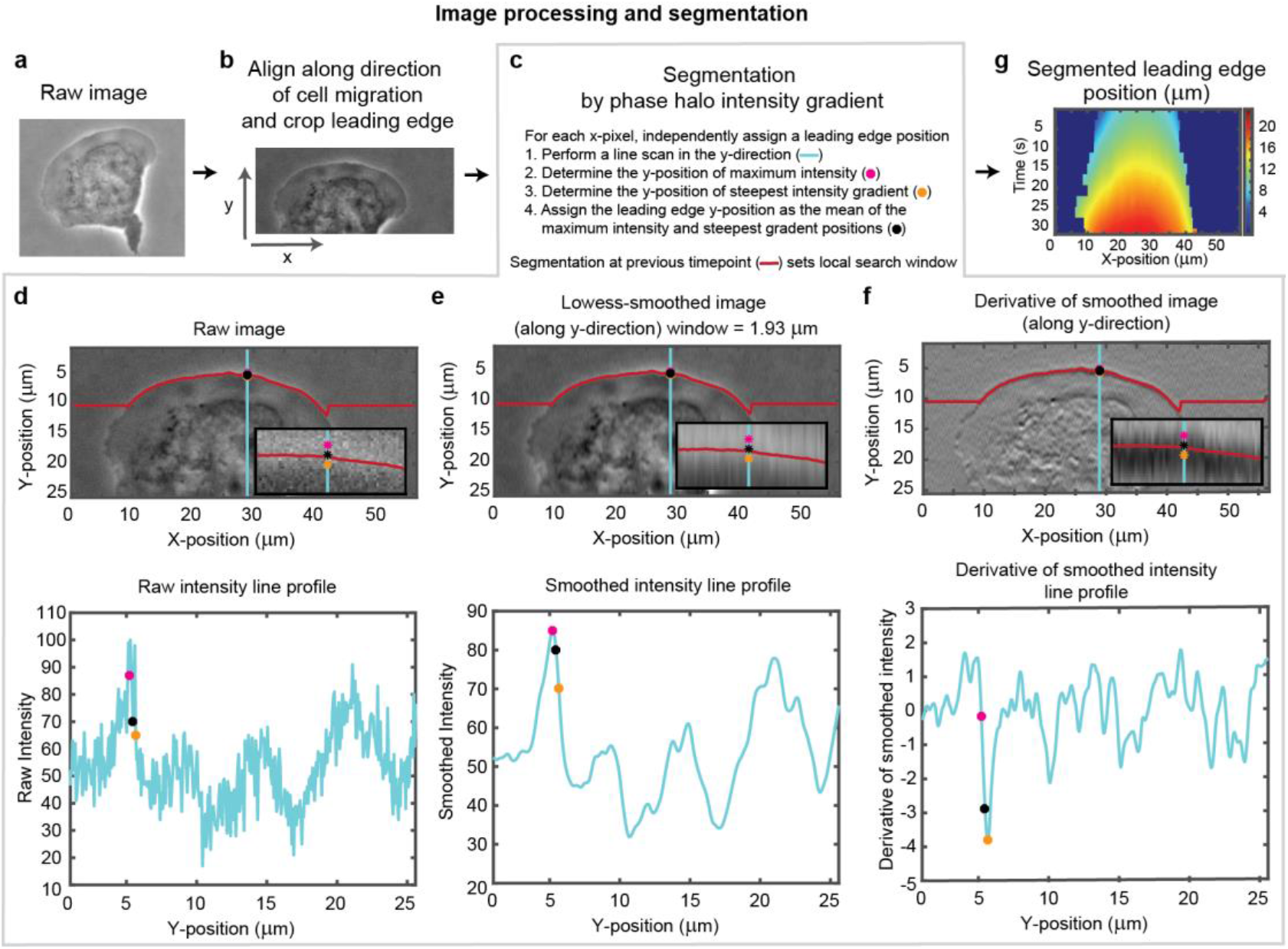
Overview of cell segmentation analysis pipeline. Example of image processing and leading edge segmentation for a single cell. (**a**) Example raw image of a migrating cell. (**b**) Image of the same cell after aligning the image in the direction of motion of the cell and then cropping the leading edge. (**c-f**) Segmentation process shown for the example cell, for a single time point. (c) Steps for performing the segmentation, as well as a legend for (d-f). (d-f) Image of the cell (top) and a vertical line scan of the pixel intensity (blue line, bottom) for (d) the raw image, (e) the image smoothed along the vertical direction, and (f) the vertical derivative of the smoothed image. Super-imposed on each plot is the position of maximum intensity (pink dot), the position of steepest negative gradient (orange dot), and the mean of these two positions (black dot), which was used as the segmented leading edge point. The search for these maxima along each line scan was performed in a window around the segmented leading edge position at the previous time point (red line). (**g**) Kymograph of the segmented leading edge positions for all time points in the video.

**Fig. S2:**
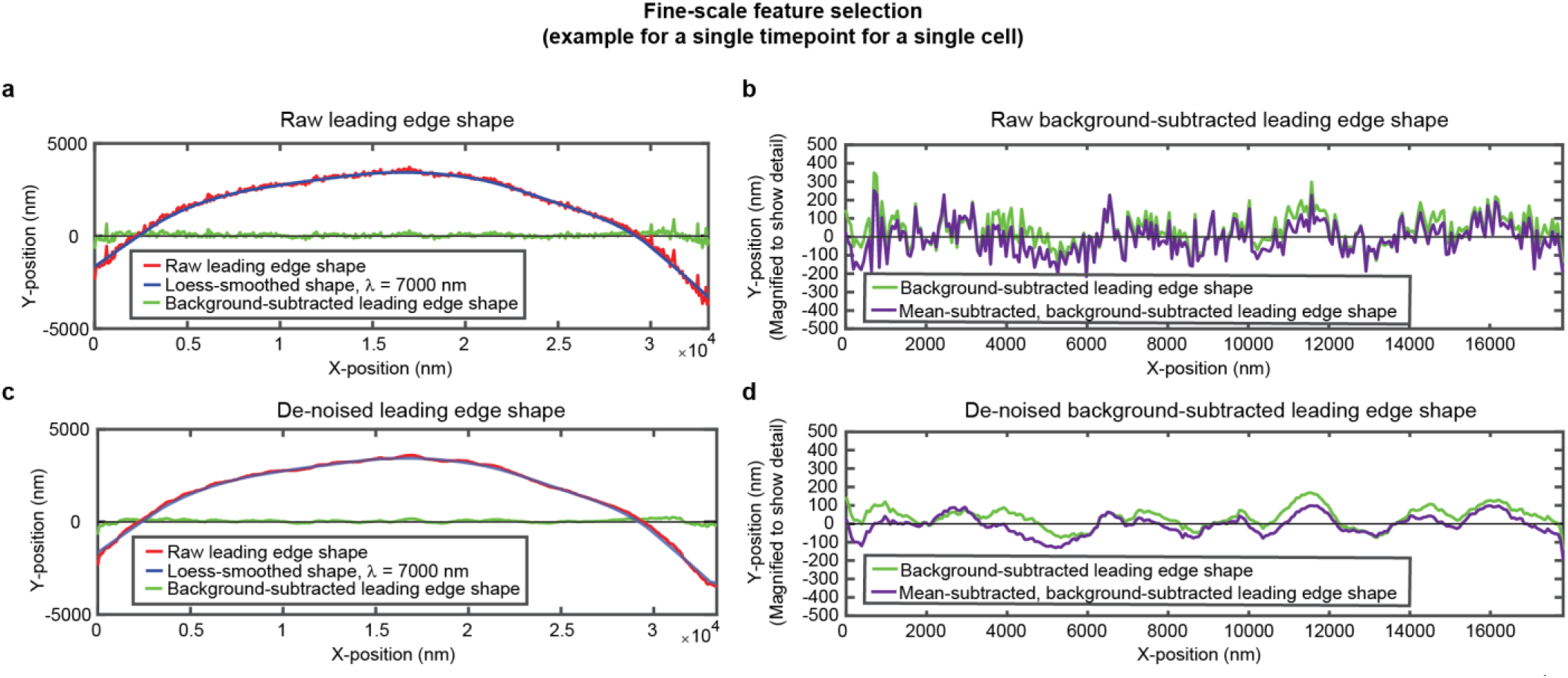
Overview of analysis pipeline to extract fine-scale leading edge shape features. An example of the fine-scale feature selection process for the same cell shown in Fig. S1. (**a,c**) Raw leading edge shape (red line), the Loess-smoothed shape (blue line), and the difference between the raw and smoothed shape (green line) for both (a) raw data and (c) de-noised data, to emphasize the ability of background-subtraction to capture fine-scale fluctuation events. (**b,d**) Green line: The same data as shown in (a,c), with the y-position magnified to show fine-scale shape features. Purple line: The same data after subtracting the time-averaged y-position for each x-position. This subtraction pulls out fluctuations around the average y-position for each x-position. The raw background-subtracted leading edge shapes (panel (b), purple curve) were used in subsequent analysis.

**Fig. S3:**
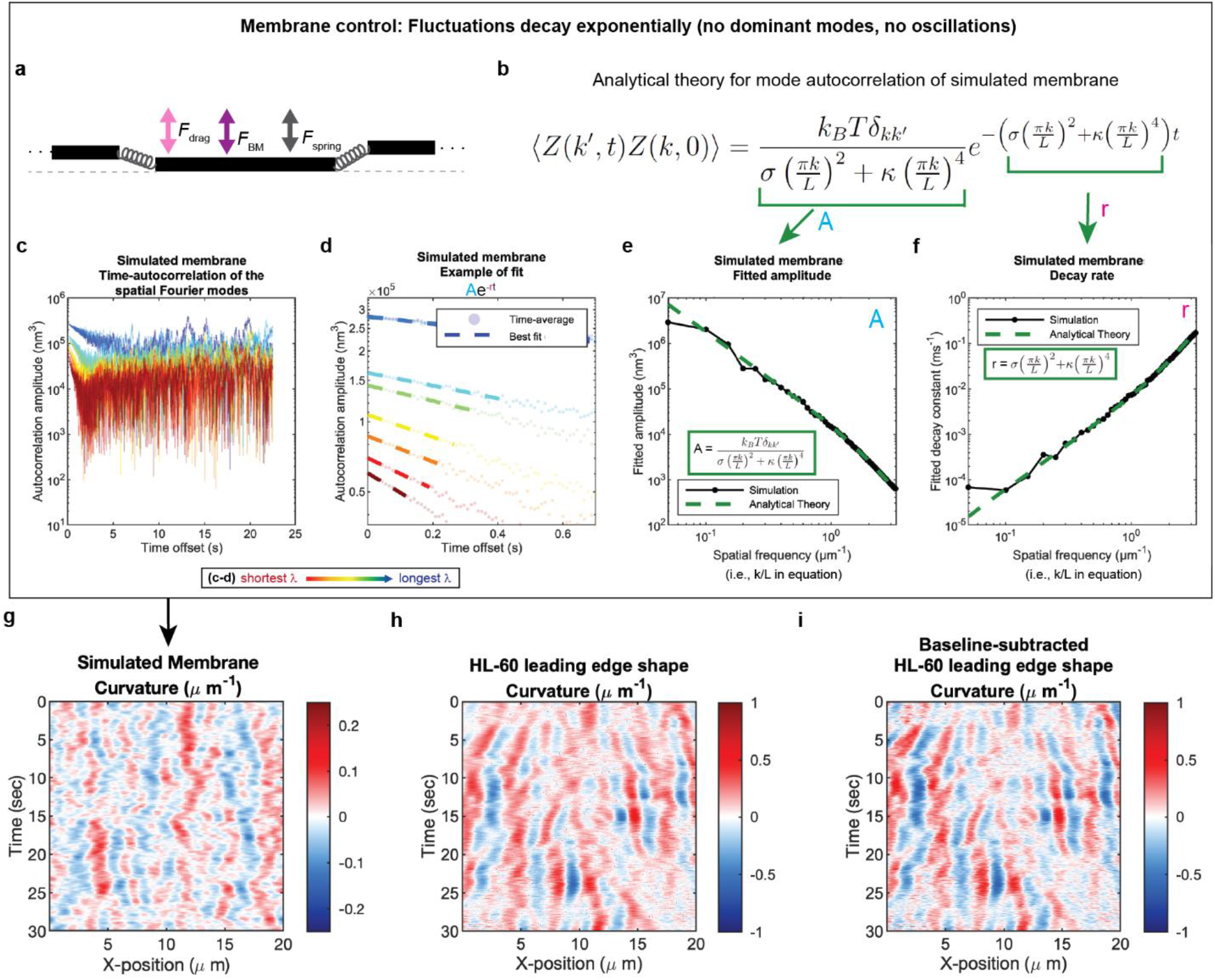
Control for Analysis I. Validation of spatial Fourier mode autocorrelation analysis using analytical theory for simulated membrane dynamics. (**a-f**) Simulated membrane control exhibiting exponentially decaying fluctuations. (a) Freely fluctuating membrane model schematic. Black lines, membrane; Physical parameters: *F*_spring_, forces between membrane segments; *F*_BM_, Brownian forces; *F*_drag_, viscous drag. (b) Analytical theory for membrane freely fluctuating under Brownian motion: Autocorrelation amplitude <*Z*(*k*’,*t*)*Z*(*k*,0)> as a function of wavemode, *k*, and time, *t*, is an exponential decay function. Parameters include the Boltzmann constant *k_B_*, temperature *T*, elastic modulus *σ*, bending modulus *κ*, and membrane rest length *L*. (**c**) Autocorrelation amplitude (complex magnitude) of the spatial Fourier transform plotted as a function of time offset from a representative simulation. Each line corresponds to a different spatial frequency in the range of 0.2-0.55 μm^-1^ (corresponding to a wavelength in the range of 5.0-1.8 μm) in 0.05 μm^-1^ intervals. (**d**) Best fit of the autocorrelation data shown in (c) to an exponential decay, fitted out to a drop in amplitude of 2/*e*. (**e-f**) Fitted parameters for the same simulation shown in (c-d) showing good agreement with the analytical theory. (**g-i**) The most obvious features of the curvature kymograph (apparent dominant wavemodes and apparent oscillations) are misleading, and are recovered for a system with (by definition) no dominant or oscillatory modes. In the experimental data, these features are retained following baseline subtraction in the pre-processing step (see Fig. S2). Curvature kymograph for (g) a simulated membrane, (h) the HL-60 leading edge displayed in Fig. 1, and (i) the same HL-60 cell leading edge shown in (h) with curvature fitting performed after baseline-subtraction.

**Fig. S4:**
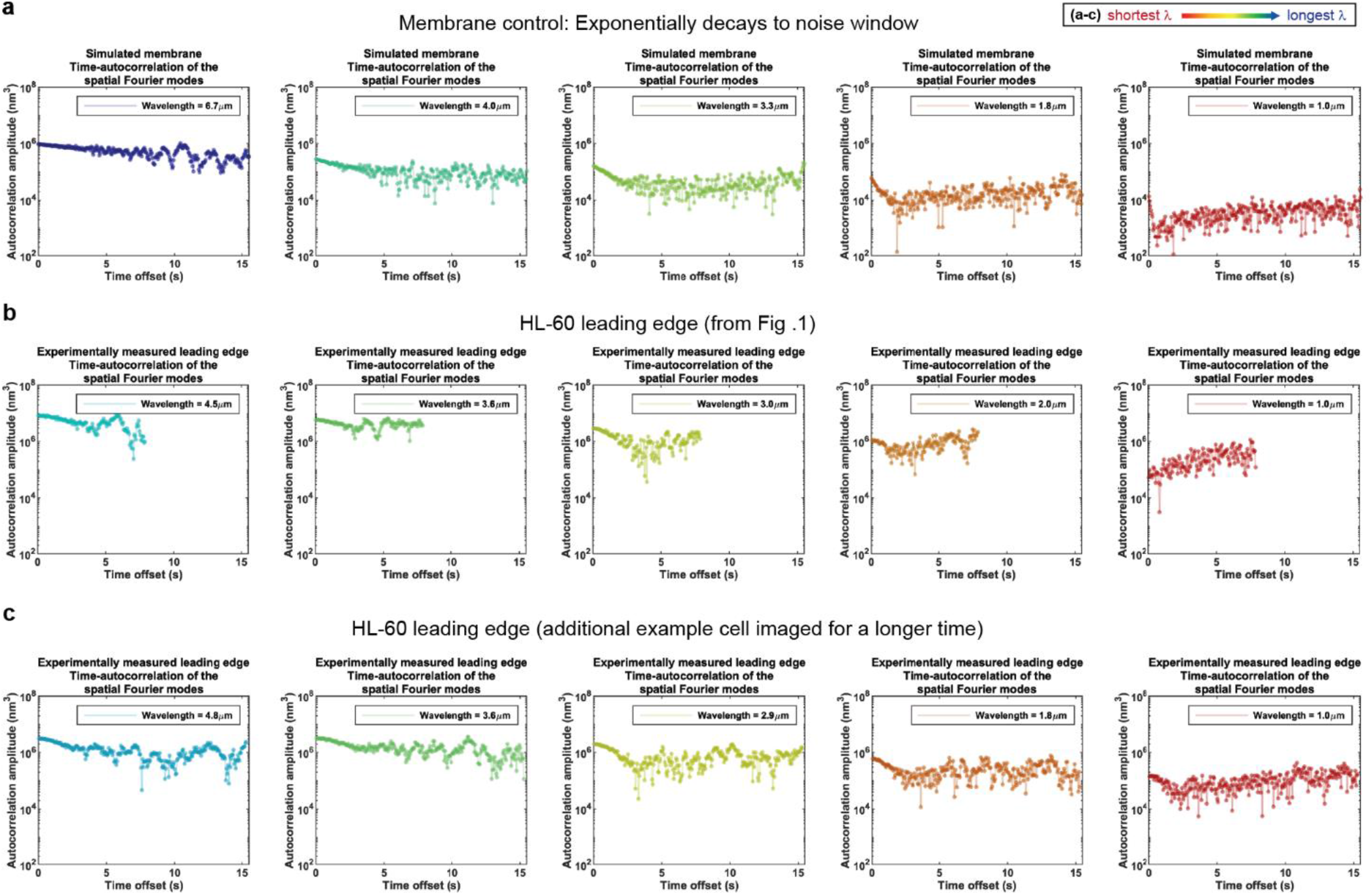
Control for Analysis II. Autocorrelation analysis of HL-60 leading edges shows exponential decay to a noise window at long times. (**a-c**) Autocorrelation amplitude (complex magnitude) of the spatial Fourier transform plotted as a function of time offset from (a) the simulated membrane control from Fig. A1, (b) the HL-60 leading edge displayed in Fig. 1, and (c) a different HL-60 cell leading edge that was imaged for longer time. Each panel shows the autocorrelation results for a different wavelength, where the color represents the wavelength.

**Fig. S5:**
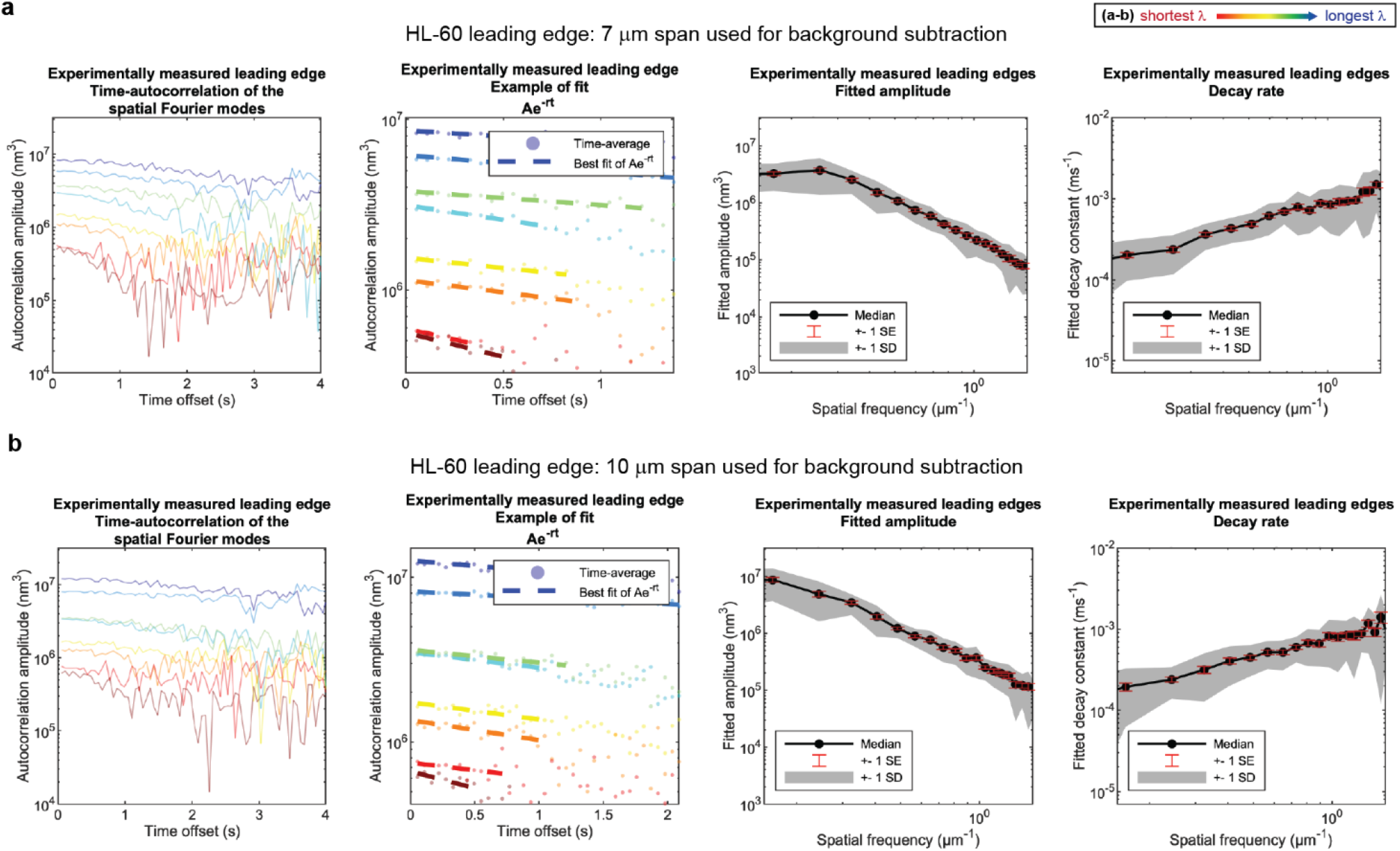
Control for Analysis III. No new features emerge upon a ~50% increase in span used for background subtraction. (**a-b**) Autocorrelation analysis results on HL-60 cell leading edges as shown in Fig. 1, using either (a) a 7 μm span or (b) a 10 μm span to perform background subtraction on the leading edge shape during pre-processing. (A typical cell is ~15-20 μm wide.)

**Fig. S6:**
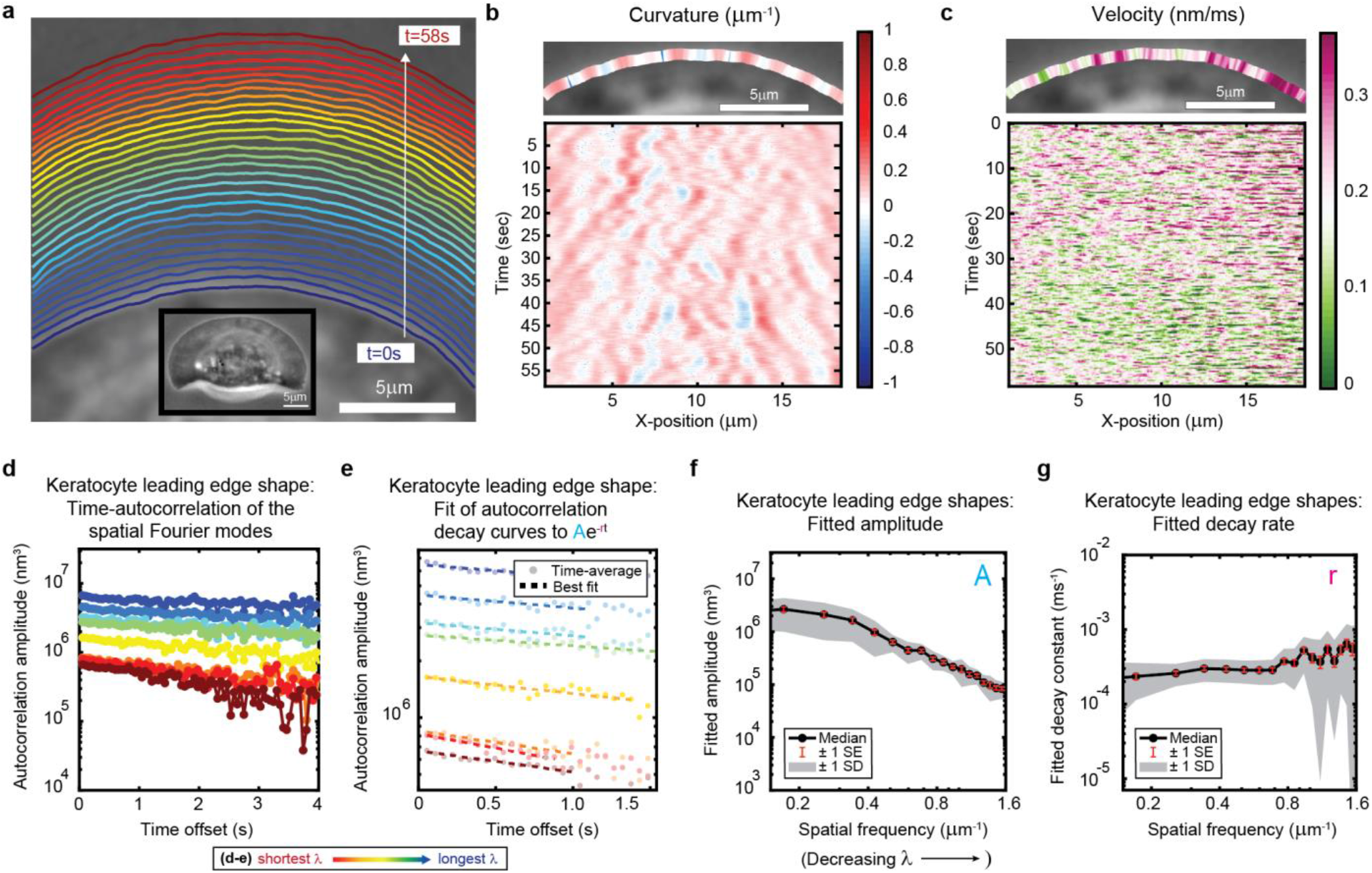
Leading edge fluctuation behavior is reproduced in fish epidermal keratocytes. (**a-c**) Example of leading edge fluctuations extracted from a representative migrating fish epidermal keratocyte, plotted as in Fig. 1a-c. Note differences in scale for time, x-position, and velocity from the equivalent plots in Fig. 1. Segmented leading edges in (a) are still plotted in 2 sec time intervals. (**d**) Autocorrelation amplitude (complex magnitude) of the spatial Fourier transform plotted as a function of time offset from a representative cell, plotted as in Fig. 1e. Each line corresponds to a different spatial frequency in the range of 0.19-0.52 μm^-1^ (corresponding to a wavelength in the range of 5.3-1.9 μm) in 0.047 μm^-1^ intervals. (**e**) Best fit of the autocorrelation data shown in (d) to an exponential decay, plotted as in Fig. 1f. Note differences in scale for time offset and autocorrelation amplitude from the equivalent plot in Fig. 1. (**f-g**) Fitted parameters of the autocorrelation averaged over 16 videos of 12 cells, plotted as in Fig. 1g-h.

**Fig. S7:**
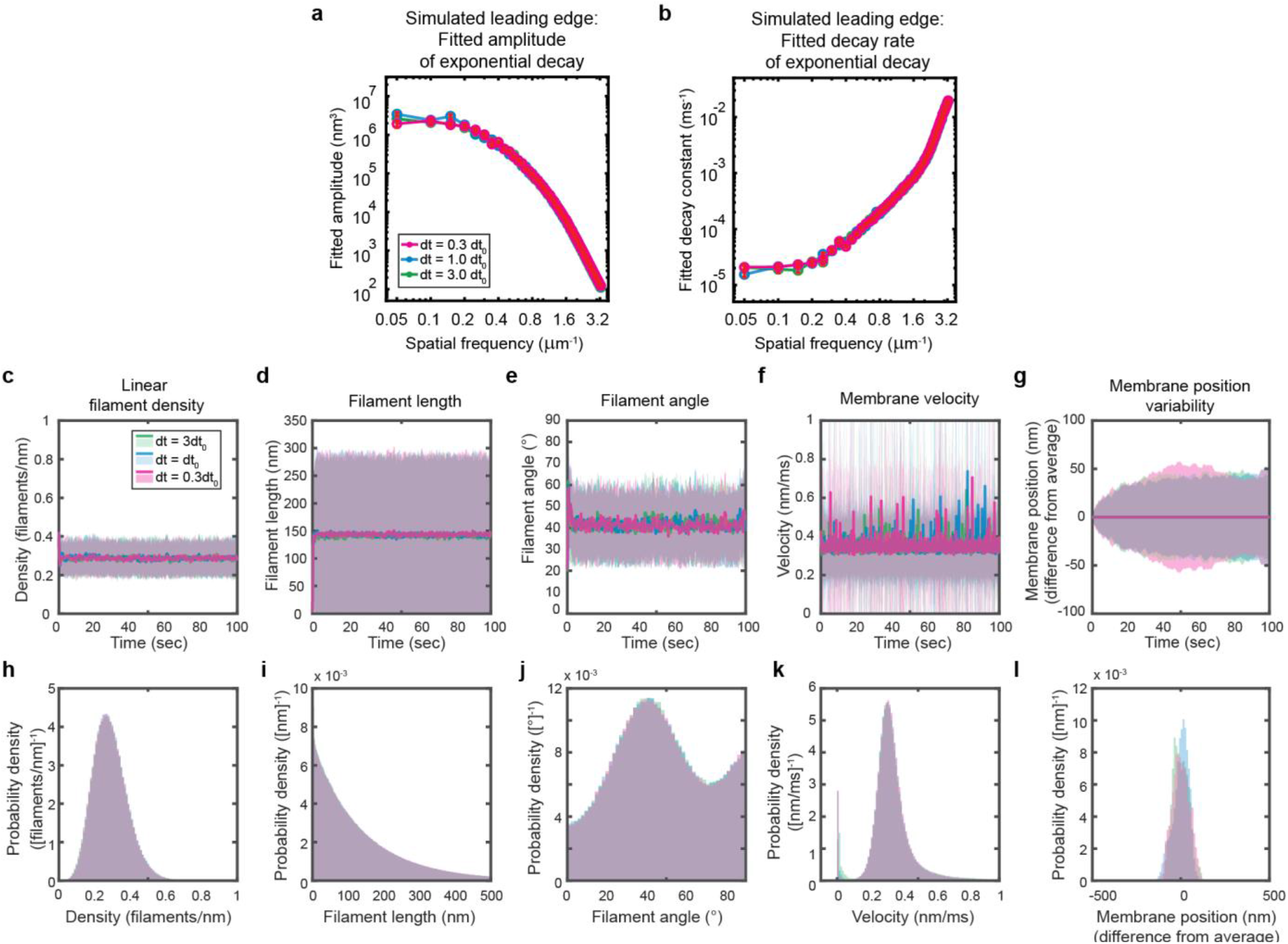
Simulations are performed at sufficient temporal discretization. A summary of leading edge fluctuations and actin network properties for simulations using the standard timestep (blue), a timestep three-fold larger (green), and a timestep three times smaller (pink). Increasing the temporal discretization (decreasing the timestep) did not affect the simulation results, suggesting this chosen timestep is appropriate to resolve the fastest timescales in the system. (**a-b**) Fitted leading edge fluctuation parameters plotted as in Fig. 5f-g. (**c-g**) Actin network properties plotted as in Fig. 2d-h. (**h-l**) Histograms of actin network property distributions at steady state (last 90 sec of the simulated timepoints in panels c-g). The y-axis corresponds to the probability density per histogram bin width. The sum of the probability density across all bins is equal to one. Histograms with bin widths less than one may have a probability density greater than one.

**Fig. S8:**
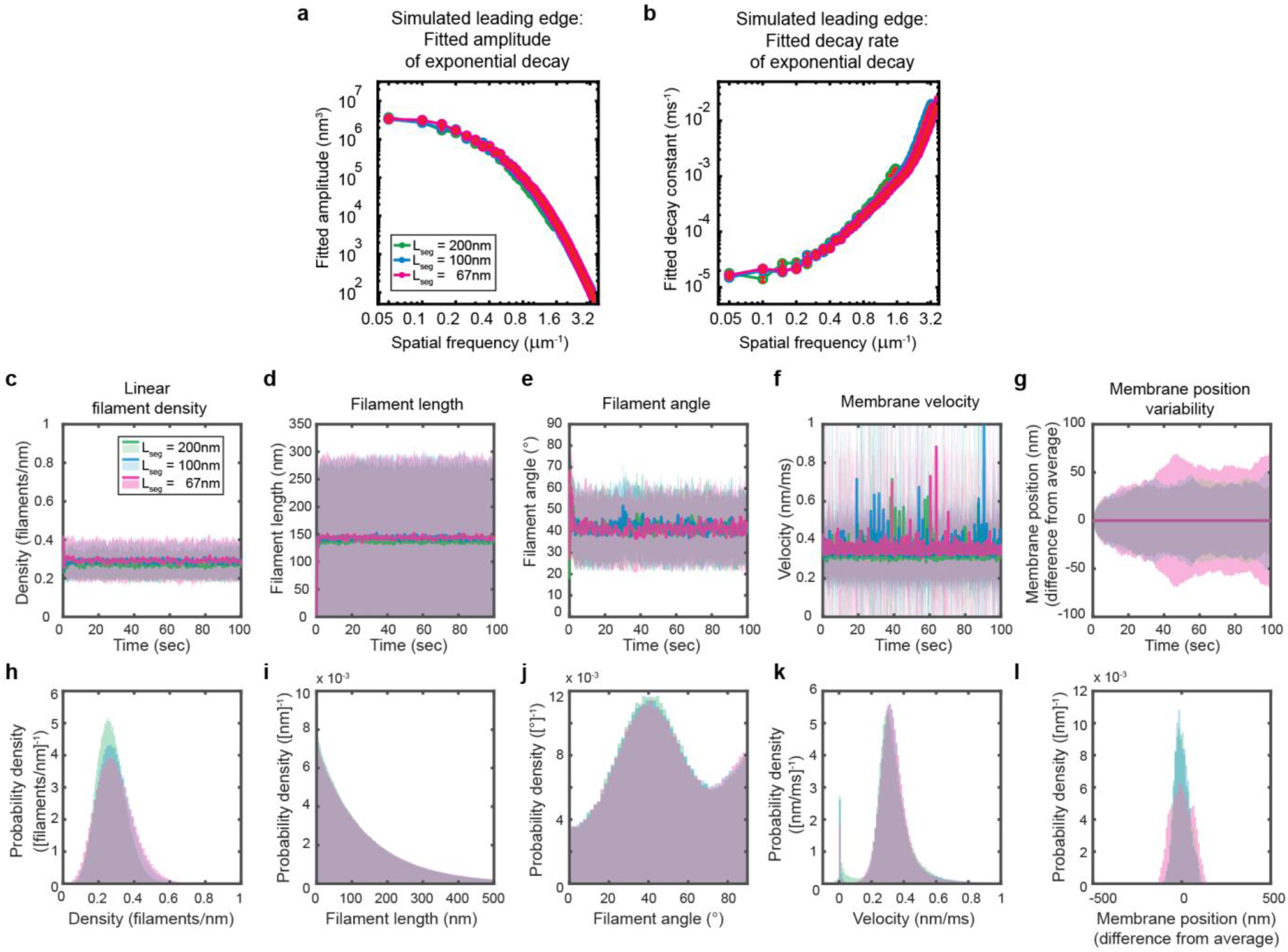
Simulations are performed at sufficient spatial discretization. (**a-l**) A summary of leading edge fluctuations and actin network properties, plotted as in Fig. S7, for simulations using the standard membrane segment length (blue), a segment size two-fold larger (green) and a length thirty percent smaller (pink). Increasing the spatial discretization (decreasing the segment size) did not significantly change the leading edge properties, suggesting the chosen discretization is sufficient to approximate a continuous membrane.

**Fig. S9:**
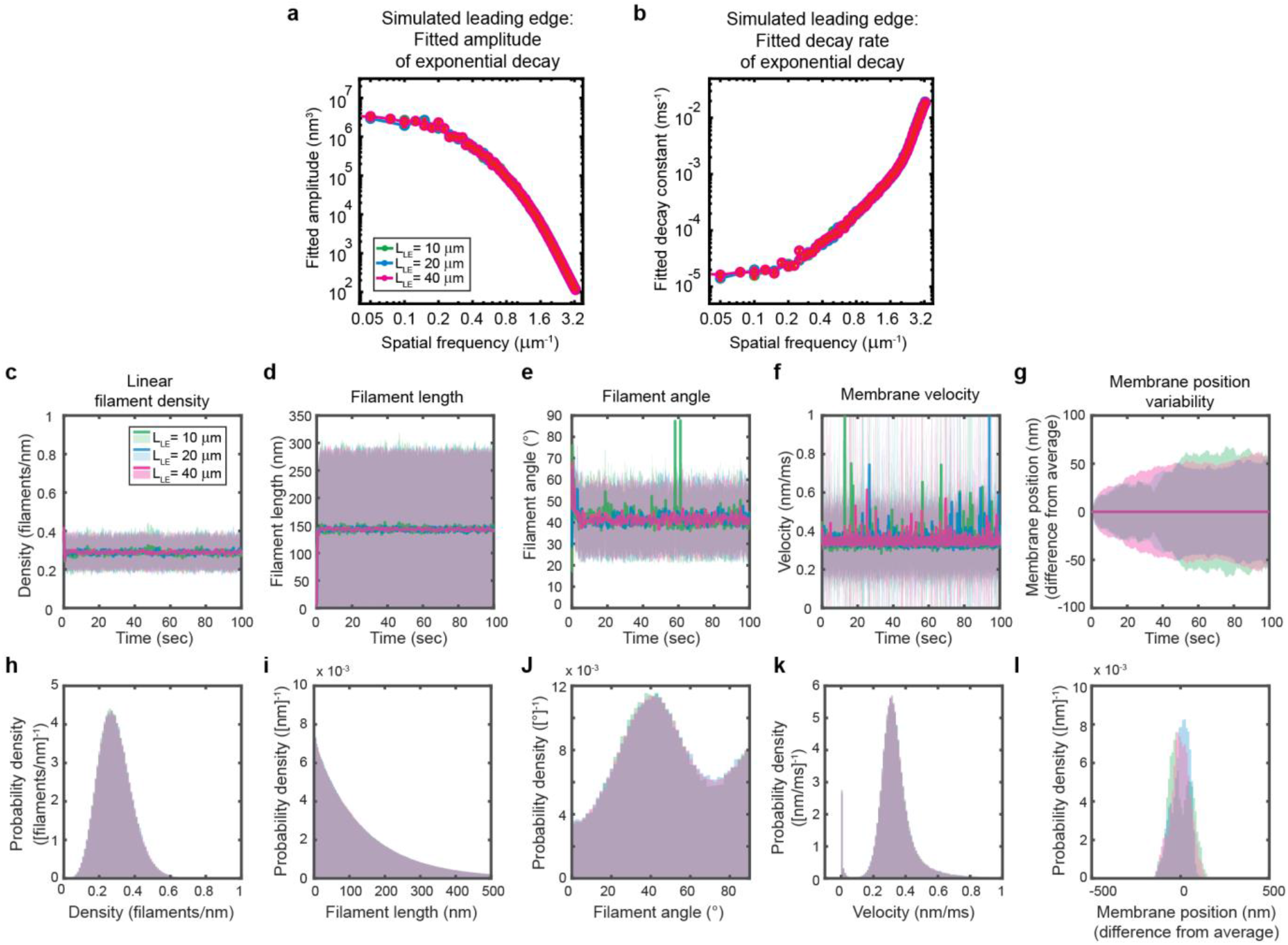
Simulated leading edge behavior is not affected by leading edge length. (**a-l**) A summary of leading edge fluctuations and actin network properties, plotted as in Fig. S7, for simulations using the standard leading edge length (blue), a length two times smaller (green) and a length two-fold larger (pink). Increasing the leading edge length did not change the properties of the fluctuations and actin network.

**Movie S1. Segmentation overlaid onto migrating HL-60 cell.** Time lapse video representation of segmentation results shown in Fig. 1a.

**Movie S2. Example fish epidermal keratocyte.** Time lapse video corresponding to the data shown in Fig. S6a-e.

**Movie S3. Example simulation.** Time lapse video representation of simulation results shown in Fig. 2c.

**Movie S4. Example HL-60 cell treated with 30nM latrunculin B.**

**Movie S5. Example HL-60 cell treated with 0.1% DMSO vehicle control.**

**Movie S6. Example HL-60 cell treated with 100 μM CK-666.**

## Appendix

### 1 Detailed description of the model

In this section, we outline the main features of the model and reference the Detailed Derivations for more detailed derivations.

#### 1.1 Model geometry

Given the flat structure of lamellipodial protrusions, we considered leading edge dynamics in two dimensions. In cartesian coordinates, the cell migrates in the x-y plane along the y-axis and maintains a fixed leading edge height of 200nm along the z-axis. The leading edge membrane was modeled as a 2D strip that restricts bending and stretching in the x-y plane and is perfectly flat along the z-axis. We discretized the membrane such that a 20 *μ*m leading edge membrane was modeled as 200 flat, rectangular segments (each 100nm in length along the x-axis and 200nm in height along z-axis) whose surface normal is fixed to lie along the y-axis. The membrane is implented using a Monge parameterization, such that these membrane segments are fixed in position along the x- and z-axes and move only along the y-axis (the direction of motion of the simulated cell).

#### 1.2 Updating the membrane position

The membrane segments are assumed to move in a viscous medium at low Reynolds number, with drag force 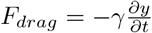 and Stoke’s drag coefficient *γ* = 6*πηr*, where r is the membrane segment length (x-axis) and *η* is the dynamic viscosity of water. The membrane also acts under the forces of membrane bending and stretching (*F_S/B_*, see Appendix section 1.3), and Brownian ratchet forces by the actin filaments (*F_BR_*, see Appendix section 1.4). This gives us the following equation of motion.

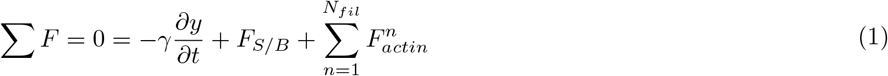

Brownian dynamics simulations, implemented with a 4th order Runge-Kutta algorithm, were performed to update the membrane segment positions. Thermal fluctuations of the membrane were ignored in this implementation, as thermal fluctuations of the (much stiffer) actin filaments dominate the membrane’s motion as well as monomer incorporation into the actin network (Mogilner and Oster 1996). See Appendix Detailed Derivations IV for a discussion of the numerical approximations included in the simulations.

#### 1.3 Forces of membrane bending and stretching

The simulated leading edge membrane acts under the energetic constrains of stretching and bending, characterized by the experimentally measurable parameters of membrane tension (*σ*, *pN* · *nm*^-1^) and bending modulus (*κ*, *pN* · *nm*), and using the following energy functional:

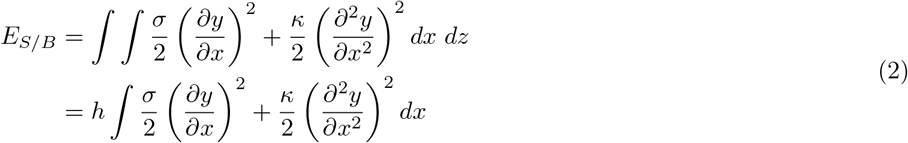

where *h* is the height of the leading edge in the z-dimension, 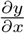 is extension of the membrane (i.e. an increase in contour length), and 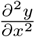 is curvature in (bending of) the membrane. The force of membrane stretching and bending on a single membrane segment (*F_S/B_*) is defined as the free energy gained by movement of the segment (*seg*), and takes the following form…(See Detailed Derivations I for a derivation of the functional derivative and resulting force.)

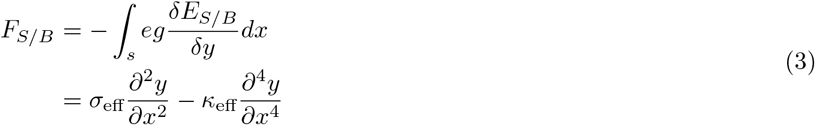

#### 1.4 Actin filament Brownian ratchet forces

Actin filaments constantly undulate due to thermal fluctuations, bending and stretching to sample their conformational space. The presence of the membrane restricts fluctuations of the filament past the membrane, presenting an entropic cost and a reduction in the free energy of the filament. The force of the filament exerted on the membrane segment, *F_actin_*, can therefore be calculated as the gain in free energy, G, by an infinitesmal movement of the membrane position, *ξ*:

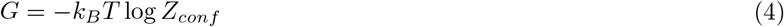

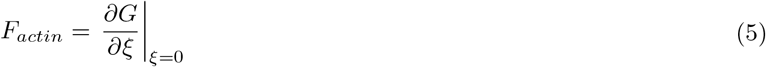

The partition function, *Z_conf_*, determined by the energetic cost of bending the filament, *E_bend_*, defines the conformational landscape. A few considerations must be made to determine the partition funciton in our system. Each filament applies force only to the membrane segment under which the filament’s barbed (growing) end equilibrium position sits, making the approximation that each membrane segment acts as an infinite wall past which filament fluctuations are blocked. Given this assumption, the membrane only restricts filament fluctuations along the y-axis. The partition function is therefore integrated over all x-positions, but only the subset of y-positions where the filament is not restricted by the membrane. With these considerations in mind, we arrive at our partition function: (See Appendix Detailed Derivations II for a derivation of the filament bending energy and Detailed Derivations III for the derivation of the force.)

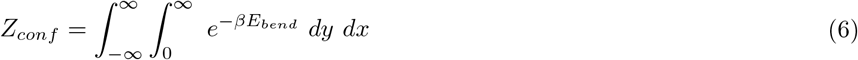

where y is measured relative to the membrane surface and 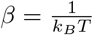. The final equation for the force of a filament on the membrane becomes…

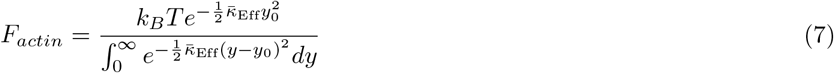

where *y*_0_ is the equilibrium filament position measured relative to the membrane surface (the filament pokes through the membrane for *y*_0_ < 0) and

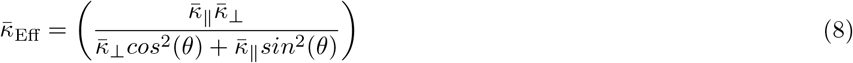

and 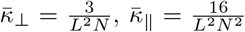, and 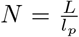 for a filament with length L and persistence length *l_p_* at an angle *θ* relative to the membrane segment normal.

#### 1.5 Actin network dynamics

Actin network dynamics including polymerization, depolymerization, branching, and capping were assumed to be independent, constant rate Poisson processes. The choice of rates are described in their respective sections below. For a given time step of Δ*t* and rate *r*, the probability of an event happening during any given time step is…

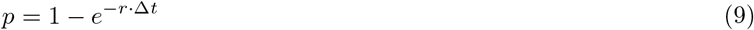

where *e*^-*r*·Δ*t*^ is the probability that the event did NOT take place within a time step of Δ*t*. A random number generator was used to determine which processes occurred in each time step. In particular, a random number was chosen between 0 and 1. If the random number lied below the probability *p*, then the event occured. For each simulation the random number generator was seeded with a unique, semi-random number, based on the current time.

##### 1.5.1 Polymerization and depolymerization

The rate of polymerization was assumed to be *r_on_* = *k_on_* · *M* · *p_gap_*, where *k_on_* is the rate of polymerization per free monomer concentration in monomers *ms*^-1^ *μM*^-1^, M is the free monomer concentration in *μM*, and *p_gap_* is the probability that enough space opens up in between the filament tip and the membrane to add a monomer (such that the polymerization rate far away from the membrane is *r*_(*y*=∞)_ = *k_on_* · *M*). Previously, it was determined that this probability is set by thermal fluctuations of the filament (Mogilner and Oster, 1996). In other words, the probability of adding a monomer is determined by the probability that thermal fluctuations, by chance, overcome the bending and stretching energies of the filament and bend the filament tip away from the membrane enough to open up a space of sufficient size to add a monomer. Given a monomer width Δ, the probability of adding a monomer is…

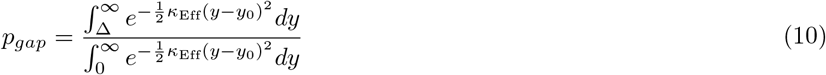

where *κ*_Eff_ takes into account the thermal energy (*k_B_T*) as well as the flexibility, length, and orientation of the filament. Depolymerization was assumed to be a constant rate process with rate *r_off_* (monomers *ms*^-1^), independent of membrane proximity.

##### 1.5.2 Branching

The rate of branching was calculated for each filament at each time step, such that *r*_branch_ = *k*_branch_ · *M* · *l*_branch_. Here *k*_branc_h is the branching rate per free monomer concentration per length of mother filament in units of branches *ms*^-1^ *μM*^-1^ *nm*^-1^, M is the free monomer concentration in *μM*, and *l*_branch_ is the length of the filament that sits inside the branching window. A fixed-length branching window is required for model stability, and is well-supported by experimental evidence that branching activation is localized to the leading edge membrane. Unlike linear polymerization, branching was not inhibited by membrane proximity. For simplicity, new branches were placed on the tip of the mother filament. If this placement caused the new branch tip position to extend past the membrane, then the branches were placed on the side of the mother filament such that the branch tip is flush with the membrane. The angle of the branch relative to the mother filament was randomly selected from a normal distribution with mean *μ* = *θ_branch_* and standard deviation *σ* = Δ*θ_branch_* as specified for each simulation. The side of the mother filament on which the branch was placed was random.

##### 1.5.3 Capping

Capping was assumed to be a constant rate process with rate *r_c_* (*ms*^-1^), independent of membrane proximity. Filaments were not allowed to uncap. Capped filaments were not allowed to polymerize or depolymerize.

##### 1.5.4 Filament deletion

Our simulations were intended to capture only leading edge actin dynamics. We therefore chose, for simulation efficiency, to only keep track of filaments actively applying force to the membrane. Filaments which were both 1) capped and 2) cumulatively provided less than 0.1% of the force on a given membrane segment were deleted from the simulation.

### 2 Detailed Derivations

This Detailed Derivations contains detailed derivations and clarifications for the material discussed in the previous section of the Appendix: “Detailed description of the model”.

#### 2.1 Detailed Derivations I: Forces of membrane bending and stretching

In this section, we use the energy functional for membrane elasticity defined in section 1.3 to calculate the elastic forces on a discrete membrane segment, filling in the steps of Appendix equation (3).

##### 2.1.1 Functional derivative of the membrane stretch/bend energy functional

We can solve the functional derivative 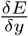 for our particular energy function (Appendix equation (2)) using the fact that for a functional of the following type…

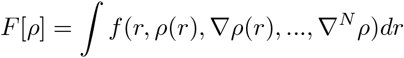

The functional derivative is calculated by…

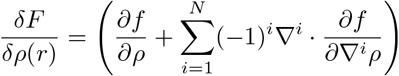

So for this particular energy functional

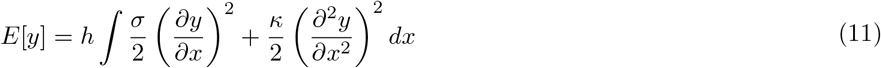

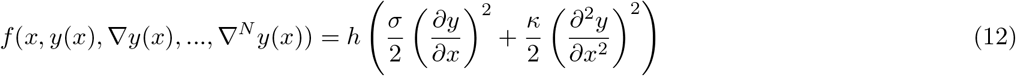

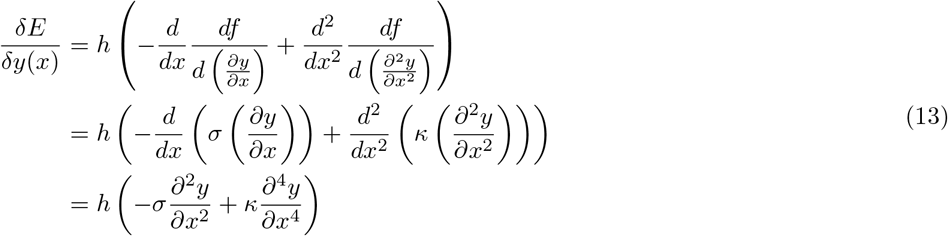

Note that 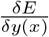 is a functional derivative (rather than an ordinary derivative), with units of a force per unit length (rather than a force). Finally, we can calculate the total elastic force on a discrete membrane segment of length Δ*x*, by integrating this force density over the length of a segment (within which the spatial derivatives of *y* do not vary).

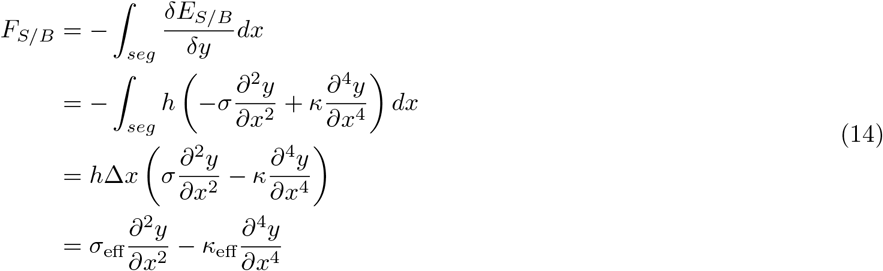

where *σ*_eff_ = *σh*Δ*x* and *κ*_eff_ = *κh*Δ*x*.

#### 2.2 Detailed Derivations II: 2D thermal fluctuations of actin filaments

In this section, we characterize the thermal fluctuations of an actin filament with a given length and persistence length (and the associated bending modulus). We first decompose the fluctuations into their (small) end-to-end and (larger) side-to-side fluctuations, and then further perform a wavelength decomposition on the side-to-side fluctuations. We next apply the equipartition theorem, giving 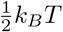 to each independent bending mode, to determine the fluctuation magnitude of each of these modes. We then determine the average total side-to-side and end-to-end fluctuation magnitudes, allowing us to calculate effective bending coefficients for the side-to-side (*κ*_⊥_) and end-to-end (*κ*_||_) fluctuations. These effective bending energies will then be used in Detailed Derivations III to determine the force of a filament on a membrane segment.

##### 2.2.1 Determining the wavelength-dependence of thermal fluctuations

Assume we have a filament of length *L*, which, in the absence of thermal fluctuations, points vertically upward in the direction 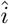. Due to thermal fluctuations, the actin polymer will fluctuate all along its length, as well as along the directions of both the short and long axis of the filament. At any point s along the polymer, we can define the local position vector 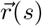, and the local tangent vector of the polymer 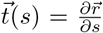. Given a persistence length *l_p_*, and thus a bending modulus *B* = *l_p_k_B_T*, the bending energy *E_bend_* of a filament is defined as…

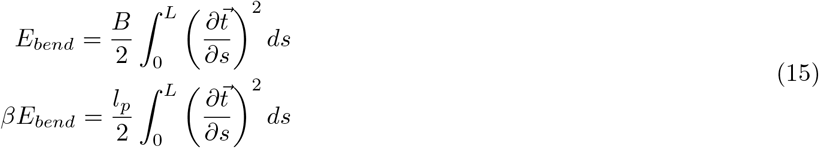

where 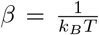 and 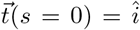. If we assume the thermal fluctuations are small 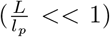, then we can write the position vector along the polymer as…

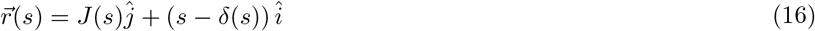

Then…

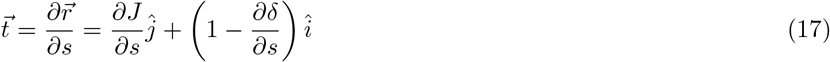

And…

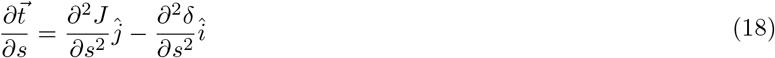

Because 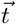 is the unit tangent vector, we know 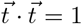. This allows us to solve for 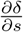 in terms of 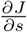.

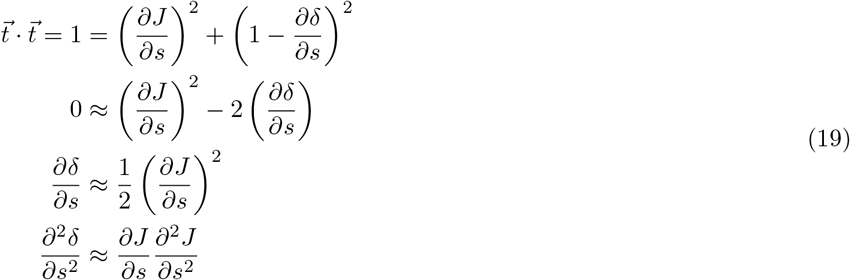

Plugging this back into our equation for *E_bend_*…

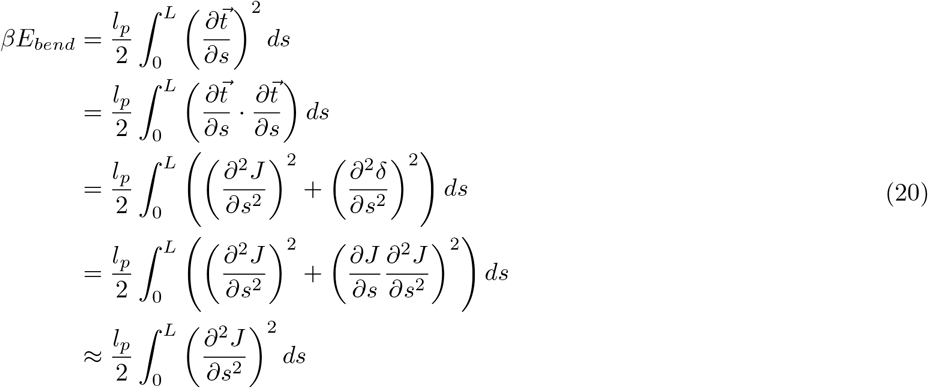

By decomposing *J*(*s*) into its wavemodes, we can determine the relative amplitudes *A_n_* of thermal fluctuations at different lengthscales, where *n* refers to the specific wavemode. This choice of wavemode decomposition oscillates around 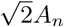 from 0 to 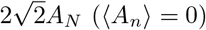, where the filament is pinned at zero at *s* = 0 and open at the other end.

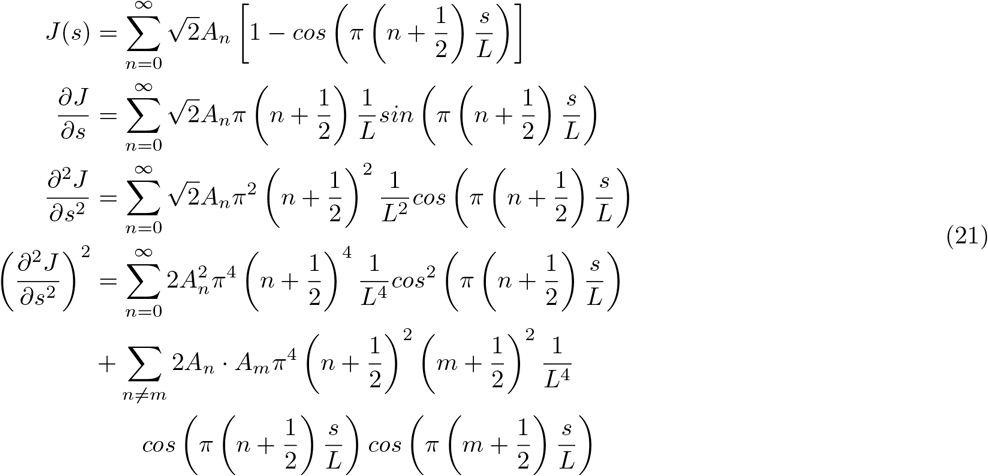

Because…

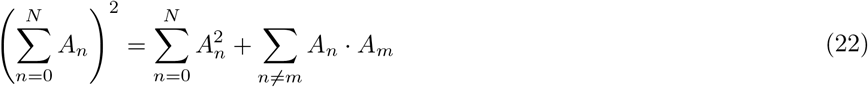

Now we can solve the integral for the parts of the function that contain *L*. For the part where *n* = *m*…

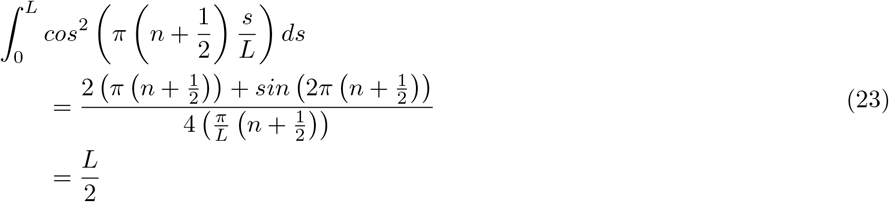

And for *n* ≠ *m*…

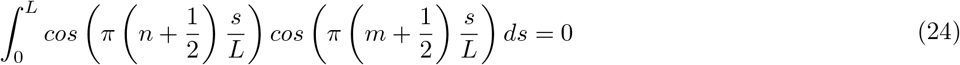

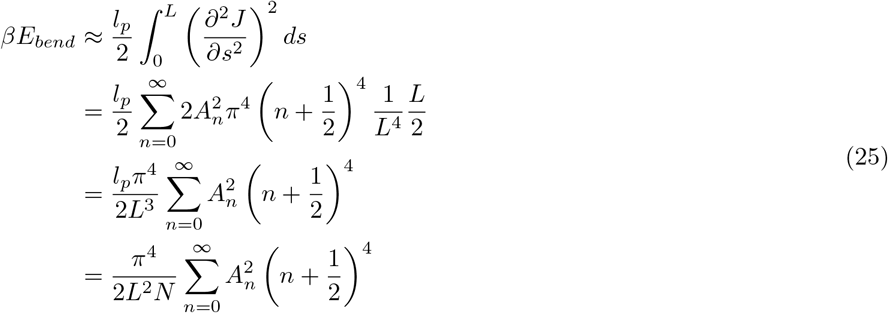

where 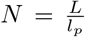. By the equipartition theorem (*βE_bend_* = 1/2 for 1 degree of freedom), we can determine the Fourier coefficients.

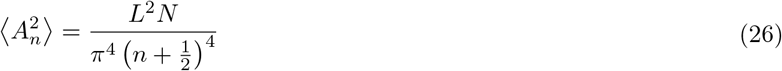

We can now calculate a few useful integrals. From the equipartition theorem and Appendix equation(26):

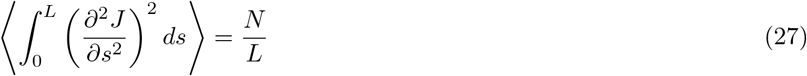

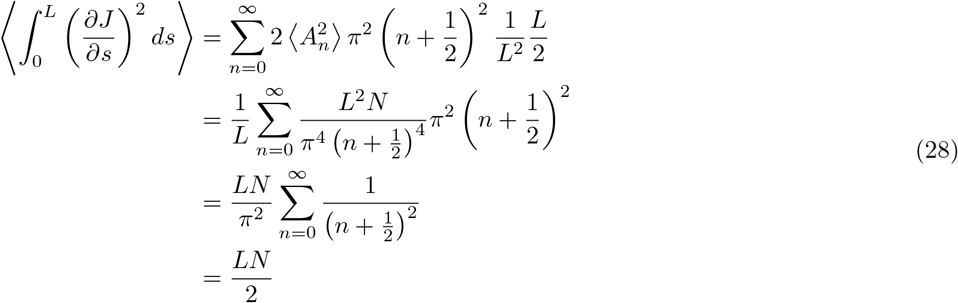

Because…

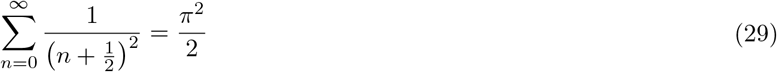

Finally…

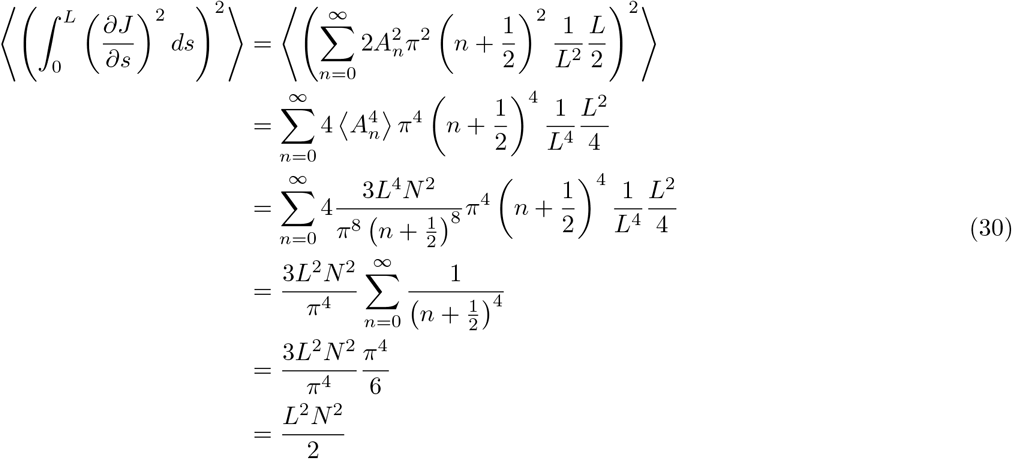

Because for a Gaussian distribution, the 4th moment is related to the variance in the following way…

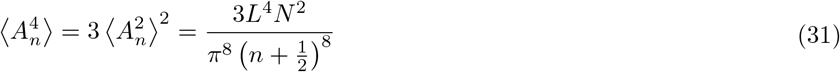

and

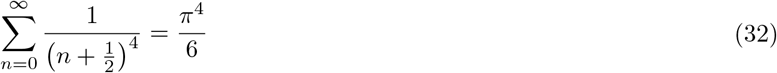

##### 2.2.2 Determining the average end-to-end retraction

The the new effective length of the filament is *I*(*s* = *L*) can be found by integrating 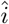 component of the tangent vector 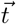 over the arc length of the filament *s*.

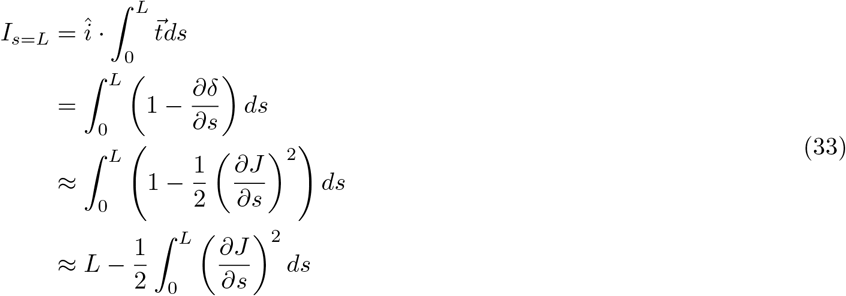

From Appendix equation(28)…

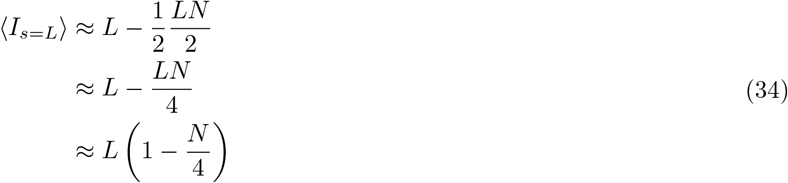

##### 2.2.3 Determining the end-to-end retraction fluctuations

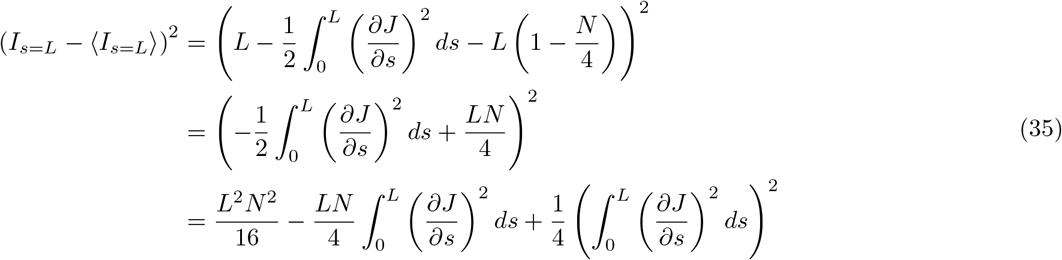

From Appendix equation(28) and equation(30)…

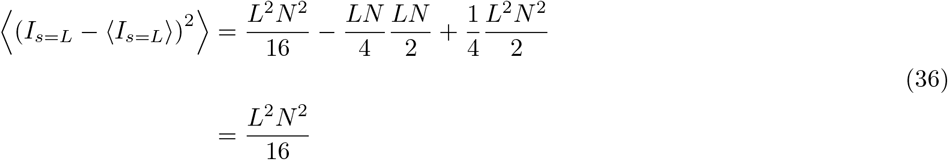

##### 2.2.4 Determining the side-to-side fluctuations

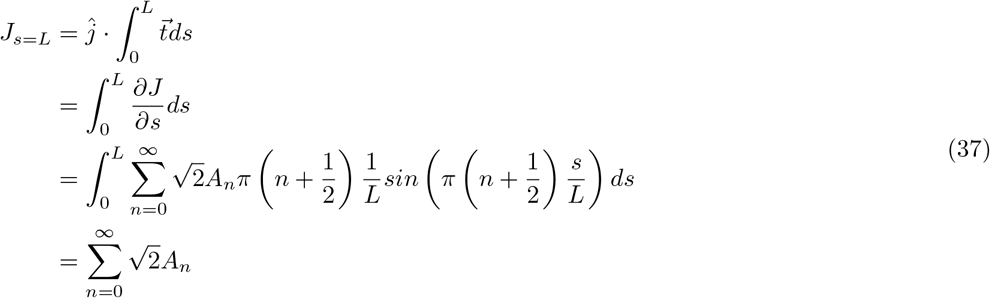

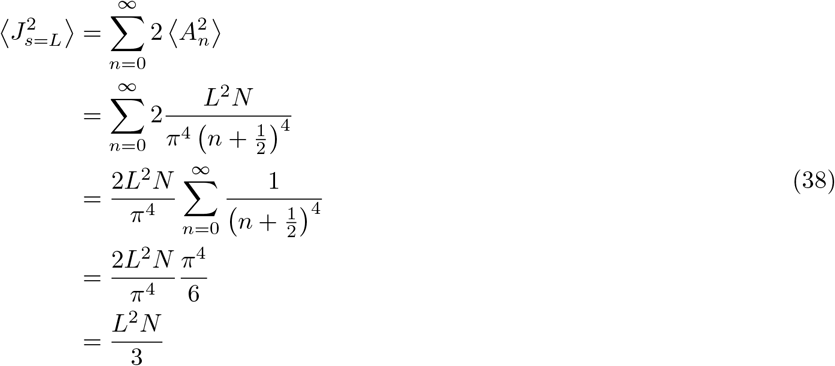

##### 2.2.5 Deriving effective stretching constants

We know by the equipartition theorem that for each degree of freedom, 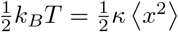. So for the 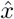 direction…

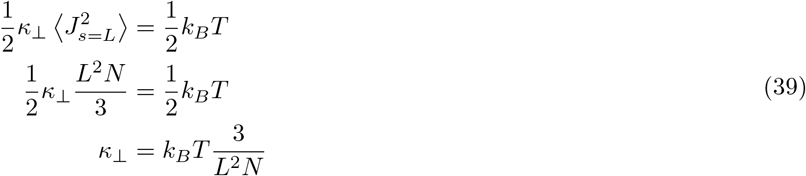

And for the 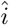 direction…

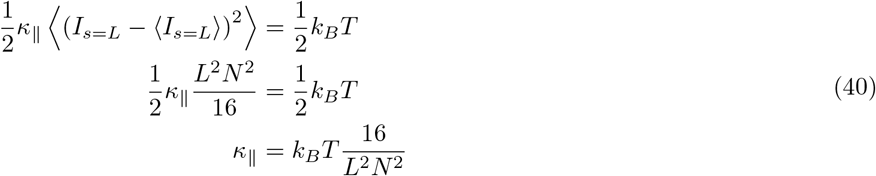

This gives us an effective bending energy as a function of actin filament tip position….

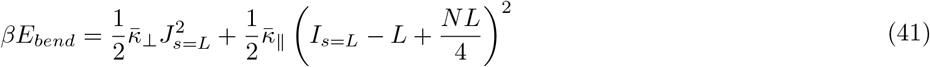

where 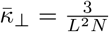 and 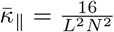.

#### 2.3 Detailed Derivations III: The force of an actin filament on the membrane

In this section, we take the effective side-to-side and end-to-end filament bending energies calculated in Detailed Derivations II to determine the force of a filament on a membrane segment. We start by converting from the coordinate system used in Detailed Derivations II (in the frame of the filament long axis) to the reference frame of the membrane segment. We then use the bending energies to evaluate the partition function *Z_conf_* for a filament which is constrained by the membrane segment – and then use the partition function to determine the free energy *G* of the filament. Finally, we evaluate the force of the filament on the membrane segment (*F_actin_*) as the increase in filament free energy obtained by an incremental movement of the membrane 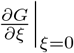.

##### 2.3.1 Converting to the reference frame of the membrane segment

We derived this bending energy function in the reference frame of the filament. However, our partition function, *Z_conf_*, will need to be integrated across the x-y coordinate system used in the rest of the paper. The energy function can be extended to an arbitrary filament orientation in a 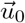 and tip position 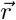 relative to resting filament tip position 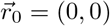 in the following way:

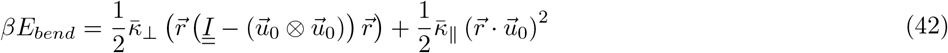

Taking a coordinate system centered around the equilibrium filament tip position and aligned with the membrane normal, a filament lying at angle *θ* relative to the membrane normal will have filament orientation vector 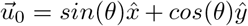 and filament tip position 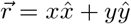 will having the following perpendicular and parallel displacements…

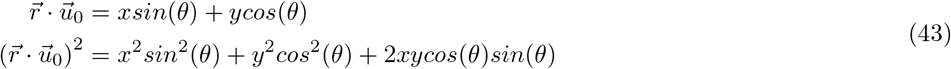

and…

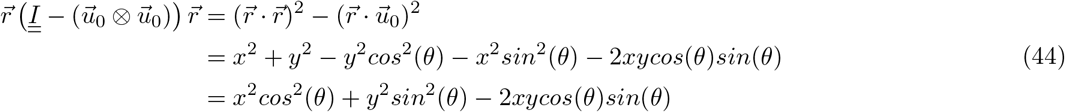

We arrive at the following bending energy as a function of the filament angle, relative to the membrane normal.

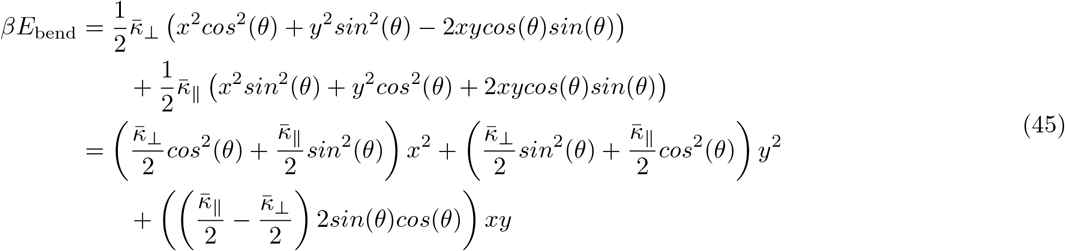

The x- and y-components have been separated for ease of integration in the next section.

##### 2.3.2 Simplification of the bending energy function

Currently, the exponential being integrated is of the form 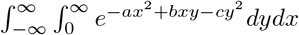. We can re-write the integral in the form 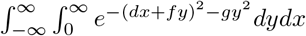, to take advantage of the fact that 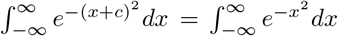. To do this, we can complete the square: 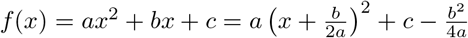

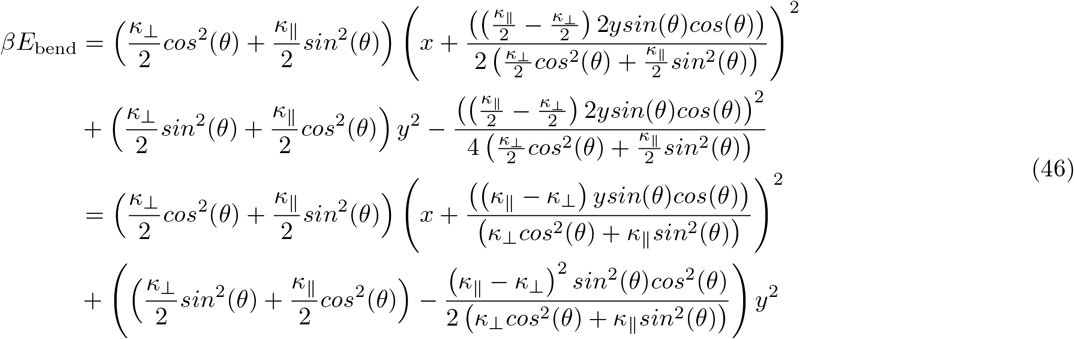

Now looking just at the *y*^2^ portion, we can determine an effective bending coefficient in the 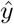 direction.

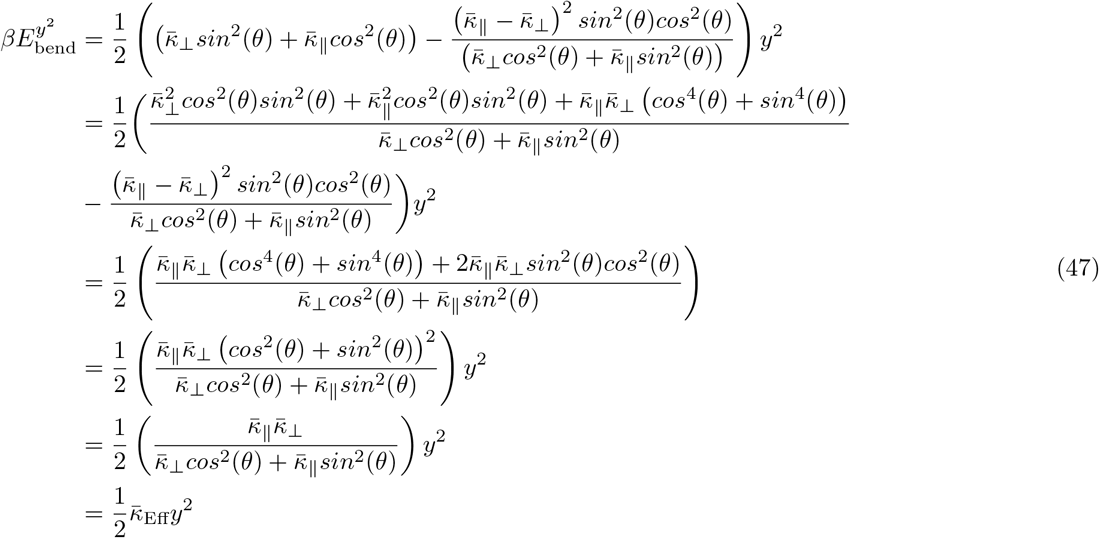

Because 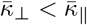 has a maximum at *θ* = 0 and a minimum at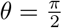. In the limit where 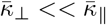, we find

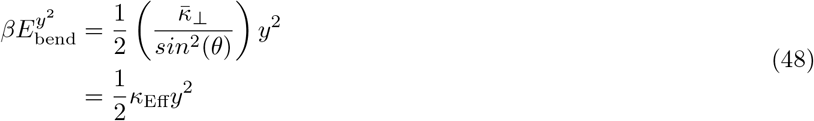

where 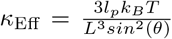, agreeing with previous models (Mogilner and Oster, 1996, BiophysJ), except for the fact that we arrive at a prefactor of 3 rather than 4, as we took into account all of the bending modes of the filament, rather than assuming bending of the filament lies along an arc of constant curvature – a difference also arrived at by Dickinson and colleagues (Dickinson, Caro, and Purich, 2004, BiophysJ).

This gives us a final bending energy…

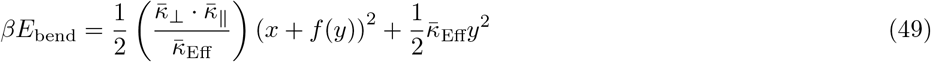

##### 2.3.3 Calculation of the partition function

Using these energies, we can determine the partition function, *Z_conf_*, summing over all possible positions of the filament. In this case, the filament can fluctuate freely along the x-axis, but cannot fluctuate past the membrane position along the y-axis. Here we define the membrane position *d* relative to the filament tip.

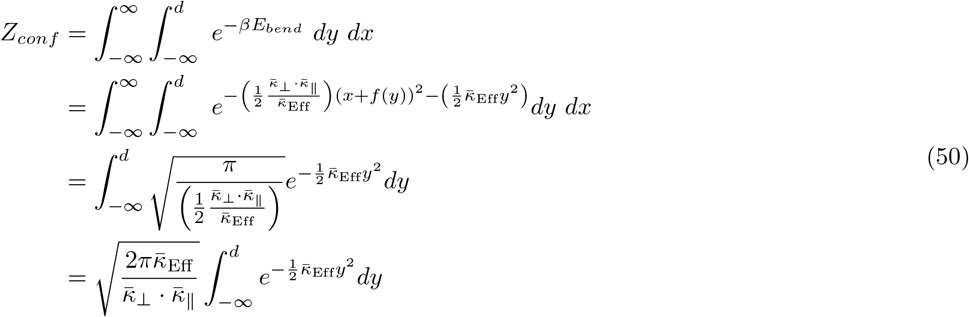

If we then have a change of variables, where we center the system at d, the equation becomes…

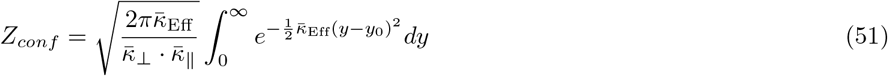

where *y*_0_ is the equilibrium filament position, measured relative to the membrane surface (the filament pokes through the membrane for *y*_0_ < 0).

##### 2.3.4 Calculation of the force of actin on a membrane

Upon an infinitesimal membrane position perturbation *ξ*, the equilibrium filament tip position becomes *y*_0_ → *y*_0_ + *ξ*. The partition function becomes…

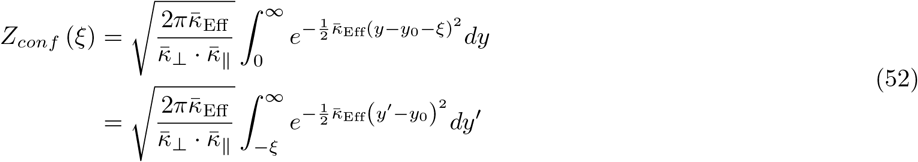

and the derivative of the partition function with respect to the perturbation is (by Leibniz’s rule)…

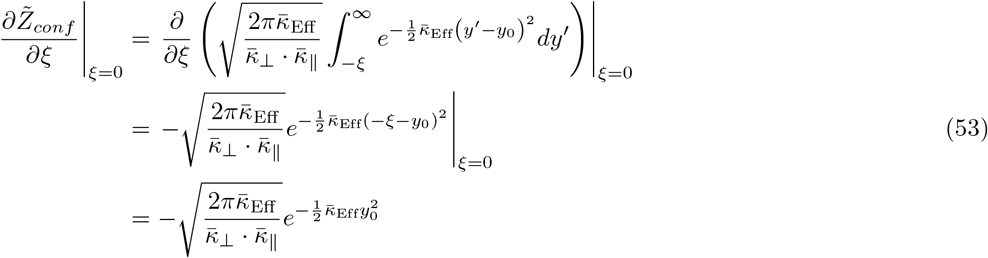

Plugging Appendix equation(51) and equation(53) into Appendix equation(5), we get the following force:

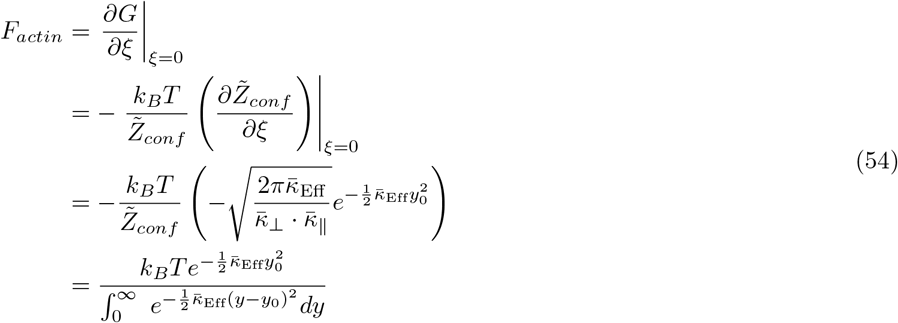

Note that this equation for the force assumes that the pointed (non-growing) ends of the filaments (as well as branches) are rigidly fixed to a stiff and immobile actin network.

#### 2.4 Detailed Derivations IV: Numerical approximations

In this section, we describe the various numerical approximations made in the development of our computational model.

##### 2.4.1 Membrane stretch/bend forces

For membrane stretch/bend force calculations, the 2nd and 4th order spatial derivatives were calculated using a central finite difference approximation with 8th order accuracy.

##### 2.4.2 Force of actin on the membrane

Many equations used in this model required performing numerical calculations of Gaussians over half-space. For this purpose, we used the error function. The denominator of Appendix equation(54) can be rewritten in terms of the error function in the following way…

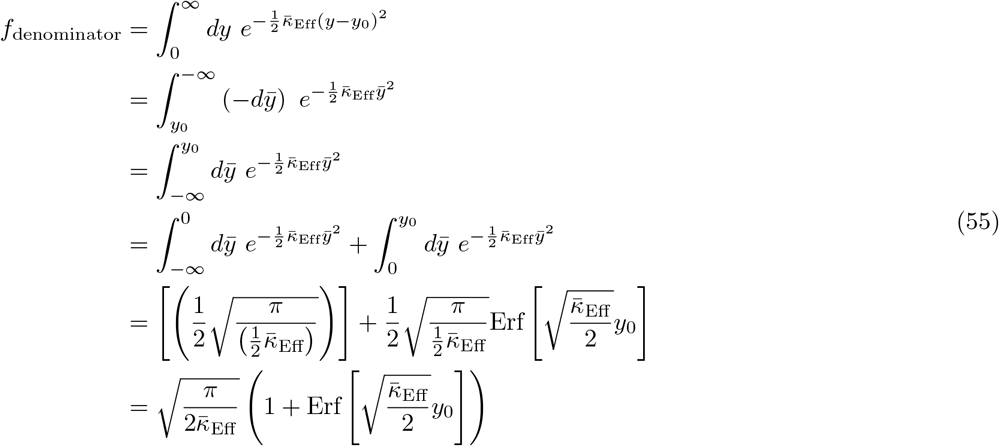

The error function appears in the equation as 1 + erf(*z*), which can give inaccurate calculations when *z* ≪ 0 and 1 + erf(*z*) ≈ 0. Therefore, in the low *z* regime (*z* < 0), we replaced 1 + erf(*z*) with erfc(|*z*|). (Because erf(-*z*) = –erf(*z*), erf(*z*(*z* < 0)) = –erf(|*z*(*z* < 0)|). It follows that 1 +erf(*z*(*z* < 0)) = 1 – erf(|*z*(*z* < 0)|) = erfc(|*z*(*z* < 0)|.) The following cases are listed below for the relevant equations.

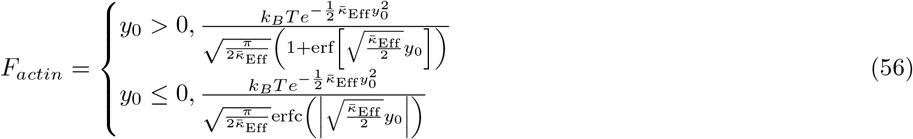

When the error function calculations failed (e.g., produced values of 0 or ∞), variable precision accuracy was used.

##### 2.4.3 Probability of adding a monomer

We used similar error function approximations to calculate the probabilities that thermal fluctuations of the filament open up a gap between the filament tip and the membrane of sufficient size to add a monomer: (See Detailed Derivations IV: Numerical approximations, Force of actin on the membrane for rational on using the erf and erfc functions.)

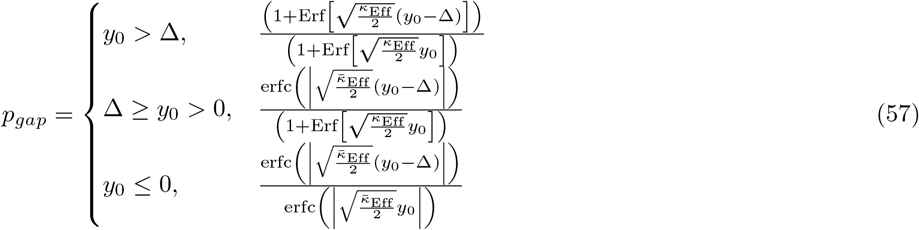

When the error function calculations failed (e.g., produced values of 0 or ∞), variable precision accuracy was used.

##### 2.4.4 Choice of timestep

The timestep was chosen to capture the fastest dynamics in the system, which could either be the membrane stretch/bend relaxation, or actin network growth dynamics. This was implemented as 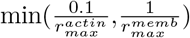. This timestep was calculated specifically for each simulation, depending on the parameters chosen. The timescales of the actin dynamics are set by the rates of polymerization, depolymerization, branching, and capping. To estimate the timescales of relaxation for membrane elastic forces, we calculate the relaxation timescales of the membrane fluctuating under Brownian thermal forces. (We do this because we do not *a priori* have an analytical theory describing the shape profile of the actin dynamics. Importantly, the fact that our simulations are not affected by the timestep (Fig. S3) provide evidence that we are sufficiently resolving all system dynamics using this approximation to set the timestep.) For a membrane at low Reynolds number, we have the following equation of motion.

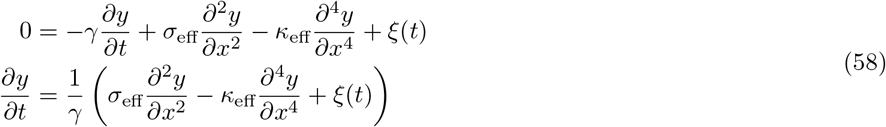

The random Brownian force is Gaussian distributed with mean *μ* = 0 and variance *σ*^2^ = 2*γk_B_T* and is defined by its autocorrelation function

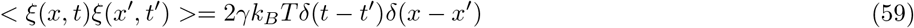

We are ultimately interested in the time evolution of wave mode solutions to this equation, so we first take the Fourier transform in order to solve for the amplitude of each wavemode. Using the following properties of the Fourier transform:

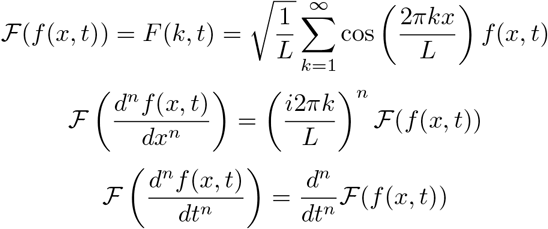

we can re-write the equation in Fourier space as…

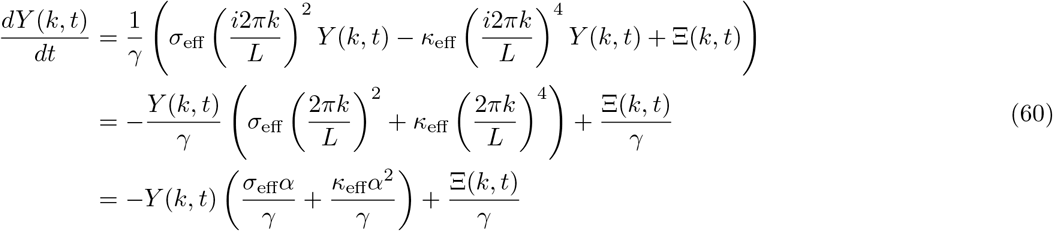

where 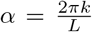. The random Brownian force in Fourier space is Gaussian distributed with mean *μ* = 0 and variance 2*γk_B_T*Δ*x* as defined by its autocorrelation function. For a discrete system, a Fourier transform with normalization 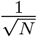, where N is the number of discrete membrane segments in this case, preserves the variance and standard deviation of a vector of normally distrbuted random values. Here, our normalization is 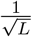 in addition to integrating over the fixed segment length Δ*x*, giving a final normalization of 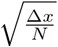 for the Brownian force 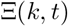. The factor of 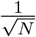 preserves the variance, leaving the variance changed only by the multiplicative factor of 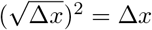.

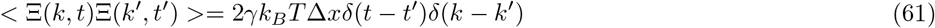

We can solve this equation using a Laplace transform.

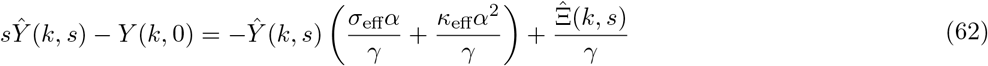

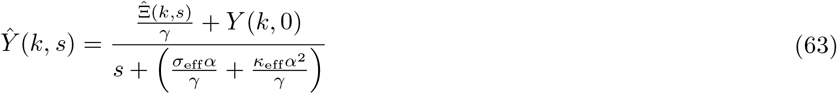

and then in inverse Laplace transform

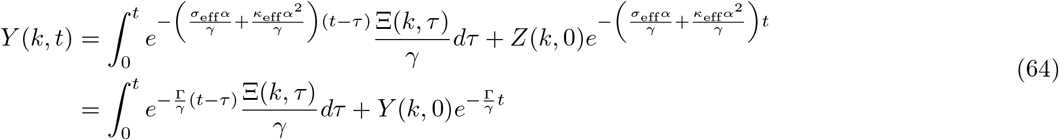

Using the properties

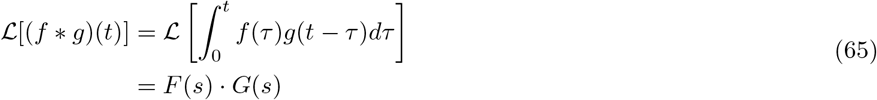

and

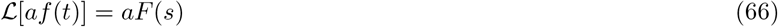

and

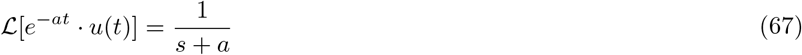

and

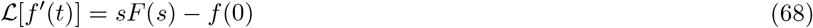

Now that we have *Y*(*k,t*), we can solve for for the time-autocorrelation function 〈*Y*(*k*’,*t*)*Y*(*k*, 0)〉.

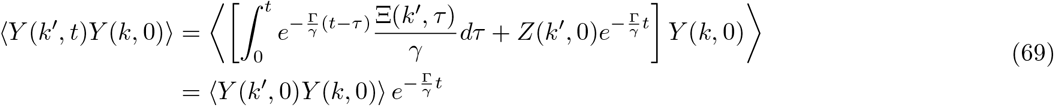

We know that 〈*Y*(*k*’, 0)*Y*(*k* 0)〉 is non-zero only if *k* = *k*’.

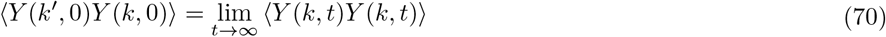

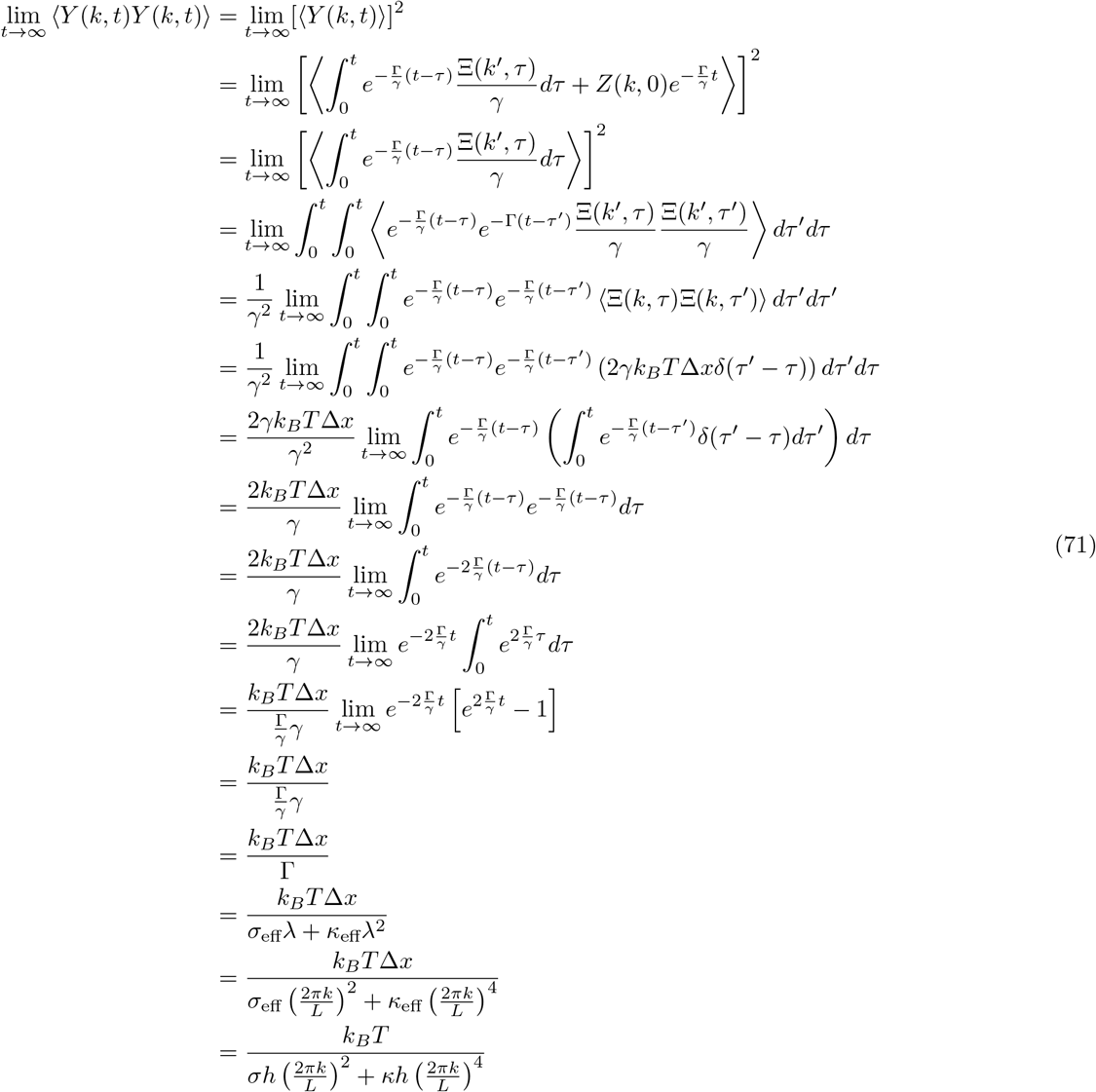

by a variant of Fubini’s theorem

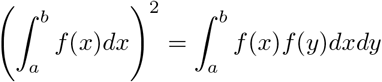

and a property of the Delta function

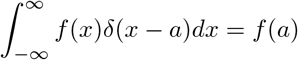

So finally…

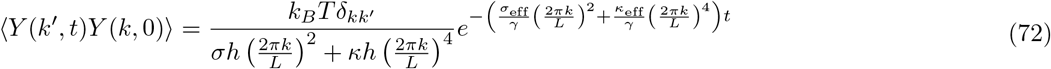

From this equation, the rate of relaxation due to membrane elasticity is…

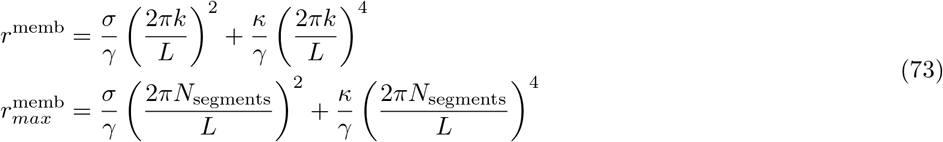

## Notes

**Competing Interests Statement:** The authors declare no competing financial interests.

https://gitlab.com/theriot_lab/leading-edge-stability-in-motile-cells-is-an-emergent-property-of-branched-actin-network-growth

### Competing Interest Statement

The authors have declared no competing interest.

## References

1. Pollard, T. D. & Cooper, J. A. Actin, a Central Player in Cell Shape and Movement. Science 326, 1208–1212 (2009).

2. Fritz-Laylin, L. K., Lord, S. J. & Mullins, R. D. WASP and SCAR are evolutionarily conserved in actin-filled pseudopod-based motility. Journal of Cell Biology 216, 1673–1688 (2017).

3. Welch, M. D., Depace, A. H., Verma, S., Iwamatsu, A. & Mitchison, T. J. The Human Arp2/3 Complex Is Composed of Evolutionarily Conserved Subunits and Is Localized to Cellular Regions of Dynamic Actin Filament Assembly. Journal of Cell Biology 138, 375–384 (1997).

4. Svitkina, T. The actin cytoskeleton and actin-based motility. Cold Spring Harbor Perspectives in Biology 10, 1–21 (2018).

5. Mogilner, A. & Oster, G. Cell motility driven by actin polymerization. Biophysical journal 71, 3030–45 (1996).

6. Peskin, C. S., Odell, G. M. & Oster, G. F. Cellular motions and thermal fluctuations: the Brownian ratchet. Biophysical Journal 65, 316–324 (1993).

7. Theriot, J. A., Mitchison, T. J., Tilney, L. G. & Portnoy, D. A. The rate of actin-based motility of intracellular Listeria monocytogenes equals the rate of actin polymerization. Nature 357, 257–260 (1992).

8. Prass, M., Jacobson, K., Mogilner, A. & Radmacher, M. Direct measurement of the lamellipodial protrusive force in a migrating cell. Journal of Cell Biology 174, 767–772 (2006).

9. Svitkina, T. M. Ultrastructure of protrusive actin filament arrays. Current Opinion in Cell Biology 25, 574–581 (2013).

10. Rafelski, S. M. & Theriot, J. A. Crawling Toward a Unified Model of Cell Motility: Spatial and Temporal Regulation of Actin Dynamics. Annual review of biochemistry 73, 209–239 (2004).

11. Vavylonis, D., Yang, Q. & O’Shaughnessy, B. Actin polymerization kinetics, cap structure, and fluctuations. Proceedings of the National Academy of Sciences of the United States of America 102, 8543–8548 (2005).

12. Svitkina, T. M., Verkhovsky, A. B., McQuade, K. M. & Borisy, G. G. Analysis of the actin-myosin II system in fish epidermal keratocytes: Mechanism of cell body translocation. Journal of Cell Biology 139, 397–415 (1997).

13. Abraham, V. C., Krishnamurthi, V., Taylor, D. L. & Lanni, F. The Actin-Based Nanomachine at the Leading Edge of Migrating Cells. Biophysical Journal 77, 1721–1732 (1999).

14. Laurent, V. M. et al. Gradient of rigidity in the lamellipodia of migrating cells revealed by atomic force microscopy. Biophysical journal 89, 667–675 (2005).

15. Fritz-Laylin, L. K. et al. Actin-based protrusions of migrating neutrophils are intrinsically lamellar and facilitate direction changes. eLife 6, (2017).

16. Tsai, T. Y. C. et al. Efficient Front-Rear Coupling in Neutrophil Chemotaxis by Dynamic Myosin II Localization. Developmental Cell 49, 189–205 (2019).

17. Lacayo, C. I. et al. Emergence of Large-Scale Cell Morphology and Movement from Local Actin Filament Growth Dynamics. PLoS Biology 5, 2035–2052 (2007).

18. De Oliveira, S., Rosowski, E. E. & Huttenlocher, A. Neutrophil migration in infection and wound repair: Going forward in reverse. Nature Reviews Immunology 16, 378–391 (2016).

19. Kolaczkowska, E. & Kubes, P. Neutrophil recruitment and function in health and inflammation. Nature Reviews Immunology 13, 159–175 (2013).

20. Diz-MuñoZ, A. et al. Membrane Tension Acts Through PLD2 and mTORC2 to Limit Actin Network Assembly During Neutrophil Migration. PLOS Biology 14, e1002474 (2016).

21. Houk, A. R. et al. Membrane Tension Maintains Cell Polarity by Confining Signals to the Leading Edge during Neutrophil Migration. Cell 148, 175–188 (2012).

22. Mueller, J. et al. Load Adaptation of Lamellipodial Actin Networks. Cell 171, 188–200 (2017).

23. Gauthier, N. C., Masters, T. A. & Sheetz, M. P. Mechanical feedback between membrane tension and dynamics. Trends in Cell Biology 22, 527–535 (2012).

24. Tsujita, K., Takenawa, T. & Itoh, T. Feedback regulation between plasma membrane tension and membrane-bending proteins organizes cell polarity during leading edge formation. Nature Cell Biology 17, 749–758 (2015).

25. Batchelder, E. L. et al. Membrane tension regulates motility by controlling lamellipodium organization. Proceedings of the National Academy of Sciences of the United States of America (2011) doi:10.1073/pnas.1010481108.

26. Sens, P. & Plastino, J. Membrane tension and cytoskeleton organization in cell motility. Journal of Physics: Condensed Matter 27, 273103 (2015).

27. Pipathsouk, A. et al. WAVE complex self-organization templates lamellipodial formation. bioRxiv (2019) doi:10.1101/836585.

28. Mullins, R. D., Bieling, P. & Fletcher, D. A. From solution to surface to filament: actin flux into branched networks. Biophysical Reviews 10, 1537–1551 (2018).

29. Risca, V. I. et al. Actin filament curvature biases branching direction. Proceedings of the National Academy of Sciences 109, 2913–2918 (2012).

30. Grimm, H. P., Verkhovsky, A. B., Mogilner, A. & Meister, J. J. Analysis of actin dynamics at the leading edge of crawling cells: implications for the shape of keratocyte lamellipodia. European Biophysics Journal 32, 563–577 (2003).

31. Maly, I. V. & Borisy, G. G. Self-organization of a propulsive actin network as an evolutionary process. Proceedings of the National Academy of Sciences 98, 11324–11329 (2001).

32. Verkhovsky, A. B. et al. Orientational Order of the Lamellipodial Actin Network as Demonstrated in Living Motile Cells. Molecular biology of the cell 14, 2559–2569 (2003).

33. Mullins, R. D., Heuser, J. A. & Pollard, T. D. The interaction of Arp2/3 complex with actin: Nucleation, high affinity pointed end capping, and formation of branching networks of filaments. Proceedings of the National Academy of Sciences 95, 6181–6186 (1998).

34. Volkmann, N. et al. Structure of arp2/3 complex in its activated state and in actin filament branch junctions. Science 293, 2456–2459 (2001).

35. Rouiller, I. et al. The structural basis of actin filament branching by the Arp2/3 complex. Journal of Cell Biology 180, 887–895 (2008).

36. Carlsson, A. E. Growth of Branched Actin Networks against Obstacles. Biophysical Journal 81, 1907–1923 (2001).

37. Parekh, S. H., Chaudhuri, O., Theriot, J. A. & Fletcher, D. A. Loading history determines the velocity of actin-network growth. Nature cell biology 7, 1219–1223 (2005).

38. Spellberg, B. J. et al. A phagocytic cell line markedly improves survival of infected neutropenic mice. Journal of Leukocyte Biology 78, 338–344 (2005).

39. Millius, A. & Weiner, O. D. Chemotaxis in Neutrophil-Like HL-60 Cells. in Chemotaxis vol. 571 167–177 (Humana Press, 2009).

40. Garner, R. M. et al. Neutrophil-like HL-60 cells expressing only GFP-tagged β-actin exhibit nearly normal motility. Cytoskeleton 77, 181–196 (2020).

41. Giannone, G. et al. Periodic lamellipodial contractions correlate with rearward actin waves. Cell 116, 431–443 (2004).

42. Ryan, G. L., Watanabe, N. & Vavylonis, D. A review of models of fluctuating protrusion and retraction patterns at the leading edge of motile cells. Cytoskeleton 69, 195–206 (2012).

43. Ma, X., Dagliyan, O., Hahn, K. M. & Danuser, G. Profiling cellular morphodynamics by spatiotemporal spectrum decomposition. PLoS Computational Biology 14, 1–29 (2018).

44. Brown, F. L. H. Elastic Modeling of Biomembranes and Lipid Bilayers. Annual Review of Physical Chemistry 59, 685–712 (2008).

45. De Gennes, P. G. Dynamics of Entangled Polymer Solutions. I. The Rouse Model. Macromolecules 9, 587–593 (1976).

46. Cerda, E. & Mahadevan, L. Geometry and Physics of Wrinkling. Physical Review Letters 90, 074302 (2003).

47. Carlsson, A. E. Growth Velocities of Branched Actin Networks. Biophysical Journal 84, 2907–2918 (2003).

48. Schaus, T. E., Taylor, E. W. & Borisy, G. G. Self-organization of actin filament orientation in the dendritic, nucleation/array-treadmilling model. Proceedings of the National Academy of Sciences of the United States of America 104, 7086–7091 (2007).

49. Schaus, T. E. & Borisy, G. G. Performance of a population of independent filaments in lamellipodial protrusion. Biophysical Journal 95, 1393–1411 (2008).

50. Bisaria, A., Hayer, A., Garbett, D., Cohen, D. & Meyer, T. Membrane-proximal F-actin restricts local membrane protrusions and directs cell migration. Science 368, 1205–1210 (2020).

51. Lieber, A. D., Yehudai-Resheff, S., Barnhart, E. L., Theriot, J. A. & Keren, K. Membrane tension in rapidly moving cells is determined by cytoskeletal forces. Current Biology 23, 1409–1417 (2013).

52. Suetsugu, S. Activation of nucleation promoting factors for directional actin filament elongation: Allosteric regulation and multimerization on the membrane. Seminars in Cell and Developmental Biology 24, 267–271 (2013).

53. Henson, J. H. et al. Arp2/3 Complex Inhibition Radically Alters Lamellipodial Actin Architecture, Suspended Cell Shape, and the Cell Spreading Process. Molecular Biology of the Cell 26, (2015).

54. Wu, C. et al. Arp2/3 is critical for lamellipodia and response to extracellular matrix cues but is dispensable for chemotaxis. Cell 148, 973–987 (2012).

55. Davidson, A. J., Amato, C., Thomason, P. A. & Insall, R. H. WASP family proteins and formins compete in pseudopod, and bleb-based migration. Journal of Cell Biology 217, 701–714 (2018).

56. Blanchoin, L. et al. Direct observation of dendritic actin filament networks nucleated by Arp2/3 complex and WASP/Scar proteins. Nature 404, 1007–1011 (2000).

57. Cai, L., Makhov, A. M., Schafer, D. A. & Bear, J. E. Coronin 1B antagonizes cortactin and remodels Arp2/3-containing actin branches in lamellipodia. Cell 134, 828–842 (2008).

58. Svitkina, T. M. & Borisy, G. G. Arp2/3 complex and actin depolymerizing factor/cofilin in dendritic organization and treadmilling of actin filament array in lamellipodia. Journal of Cell Biology 145, 1009–1026 (1999).

59. Huang, W. Y. C. et al. Phosphotyrosine-mediated LAT assembly on membranes drives kinetic bifurcation in recruitment dynamics of the Ras activator SOS. Proceedings of the National Academy of Sciences 113, 8218–8223 (2016).

60. Battich, N., Stoeger, T. & Pelkmans, L. Control of Transcript Variability in Single Mammalian Cells. Cell 163, 1596–1610 (2015).

61. Chang, A. Y. & Marshall, W. F. Organelles - Understanding noise and heterogeneity in cell biology at an intermediate scale. Journal of Cell Science 130, 819–826 (2017).

62. Mohapatra, L., Goode, B. L., Jelenkovic, P., Phillips, R. & Kondev, J. Design Principles of Length Control of Cytoskeletal Structures. Annual Review of Biophysics 45, 85–116 (2016).

63. Raj, A. & van Oudenaarden, A. Nature, Nurture, or Chance: Stochastic Gene Expression and Its Consequences. Cell 135, 216–226 (2008).

64. Gray, W. T. et al. Nucleoid Size Scaling and Intracellular Organization of Translation across Bacteria. Cell 177, 1632–1648.e20 (2019).

65. Oates, A. C. What’s all the noise about developmental stochasticity? Development 138, 601–607 (2011).

66. Welf, E. S. & Danuser, G. Using fluctuation analysis to establish causal relations between cellular events without experimental perturbation. Biophysical Journal 107, 2492–2498 (2014).

67. Mogilner, A. & Oster, G. Force generation by actin polymerization II: The elastic ratchet and tethered filaments. Biophysical Journal 84, 1591–1605 (2003).

68. Soo, F. S. & Theriot, J. A. Adhesion controls bacterial actin polymerization-based movement. Proceedings of the National Academy of Sciences 102, 16233–16238 (2005).

69. Alberts, J. B. & Odell, G. M. In silico reconstitution of Listeria propulsion exhibits nano-saltation. PLoS Biology 2, (2004).

70. Kuo, S. C. & McGrath, J. L. Steps and fluctuations of Listeria monocytogenes during actin-based motility. Nature 407, 1026–1029 (2000).

71. Chaudhuri, O., Parekh, S. H. & Fletcher, D. A. Reversible stress softening of actin networks. Nature 445, 295–298 (2007).

72. Zhao, H., Pykäläinen, A. & Lappalainen, P. I-BAR domain proteins: linking actin and plasma membrane dynamics. Current Opinion in Cell Biology 23, 14–21 (2011).

73. Devreotes, P. N. et al. Excitable Signal Transduction Networks in Directed Cell Migration. Annual Review of Cell and Developmental Biology 33, 103–125 (2017).

74. Lou, S. S., Diz-Muñoz, A., Weiner, O. D., Fletcher, D. A. & Theriot, J. A. Myosin light chain kinase regulates cell polarization independently of membrane tension or Rho kinase. Journal of Cell Biology 209, 275–288 (2015).

75. Seroussi, I., Veikherman, D., Ofer, N., Yehudai-Resheff, S. & Keren, K. Segmentation and tracking of live cells in phase-contrast images using directional gradient vector flow for snakes. Journal of Microscopy 247, 137–146 (2012).

76. Cooper, J. A. The Role of Actin Polymerization in Cell Motility. Annual Review of Physiology 53, 585–605 (1991).

77. Marchand, J. B. et al. Actin-based movement of listeria monocytogenes: Actin assembly results from the local maintenance of uncapped filament barbed ends at the bacterium surface. Journal of Cell Biology 130, 331–343 (1995).

78. Pollard, T. D. Rate constants for the reactions of ATP - and ADP-actin with the ends of actin filaments. Journal of Cell Biology 103, 2747–2754 (1986).

79. Schafer, D. A., Jennings, P. B. & Cooper, J. A. Dynamics of capping protein and actin assembly in vitro: Uncapping barbed ends by polyphosphoinositides. Journal of Cell Biology 135, 169–179 (1996).

80. Pollard, T. D., Blanchoin, L. & Mullins, R. D. Molecular Mechanisms Controlling Actin Filament Dynamics in Nonmuscle Cells. Annu. Rev. Biophys. Biomol. Struct. 29, 545–576 (2000).

81. Käs, J. et al. F-actin, a model polymer for semiflexible chains in dilute, semidilute, and liquid crystalline solutions. Biophysical Journal 70, 609–625 (1996).

82. Wirtz, D. Particle-Tracking Microrheology of Living Cells: Principles and Applications. Annual Review of Biophysics 38, 301–326 (2009).

83. Urban, E., Jacob, S., Nemethova, M., Resch, G. P. & Small, J. V. Electron tomography reveals unbranched networks of actin filaments in lamellipodia. Nature Cell Biology 12, 429–435 (2010).

